# Discovery of a druggable copper-signaling pathway that drives cell plasticity and inflammation

**DOI:** 10.1101/2022.03.29.486253

**Authors:** Stéphanie Solier, Sebastian Müller, Tatiana Cañeque, Antoine Versini, Leeroy Baron, Pierre Gestraud, Nicolas Servant, Laila Emam, Arnaud Mansart, G. Dan Pantoș, Vincent Gandon, Valentin Sencio, Cyril Robil, François Trottein, Anne-Laure Bègue, Hélène Salmon, Sylvère Durand, Ting-Di Wu, Nicolas Manel, Alain Puisieux, Mark A. Dawson, Sarah Watson, Guido Kroemer, Djillali Annane, Raphaël Rodriguez

## Abstract

Inflammation is a complex physiological process triggered in response to harmful stimuli. It involves specialized cells of the immune system able to clear sources of cell injury and damaged tissues to promote repair. Excessive inflammation can occur as a result of infections and is a hallmark of several diseases. The molecular basis underlying inflammatory responses are not fully understood. Here, we show that the cell surface marker CD44, which characterizes activated immune cells, acts as a metal transporter that promotes copper uptake. We identified a chemically reactive pool of copper(II) in mitochondria of inflammatory macrophages that catalyzes NAD(H) redox cycling by activating hydrogen peroxide. Maintenance of NAD^+^ enables metabolic and epigenetic programming towards the inflammatory state. Targeting mitochondrial copper(II) with a rationally-designed dimer of metformin triggers distinct metabolic and epigenetic states that oppose macrophage activation. This drug reduces inflammation in mouse models of bacterial and viral (SARS-CoV-2) infections, improves well-being and increases survival. Identifying mechanisms that regulate the plasticity of immune cells provides the means to develop next-generation medicine. Our work illuminates the central role of copper as a regulator of cell plasticity and unveils a new therapeutic strategy based on metabolic reprogramming and the control of epigenetic cell states.

---

Inflammation can occur as a result of infection. It is a complex biological process regulated by pro-inflammatory and anti-inflammatory mediators that orchestrate a balanced response to enable clearance of pathogens and tissue repair^1^. When this balance is lost, excessive inflammation driven by macrophages and other immune cells results in tissue injury and organ failure^2,3^. Sepsis, a deregulated host-response to infection, places a major burden on healthcare systems with approximately 50 million cases per year and 11 million deaths, which represent 1 out of 5 of global mortalities^4^. Effective drugs against severe forms of inflammation are scarce^5,6^, calling for therapeutic innovation.

Macrophages are immune cells that mediate tissue repair and host defense against pathogens, in particular through the production of growth factors and cytokines. An excessive response of macrophages to infection can be damaging^7-9^. Inflammatory macrophages are characterized by elevated expression of CD44, a cell surface marker involved in inflammation^10^. CD44-deficient mice have a reduced capacity to resolve lung inflammation and accumulate hyaluronic acids (HA) in lungs, while reconstituted CD44-positive alveolar macrophages reverse the inflammatory phenotype^11^. The molecular bases underlying the role of CD44 in inflammation remain elusive.

HA are negatively charged biopolymers that interact with positively charged metal ions. The recent finding that CD44 mediates endocytosis of iron-bound HA in cancer cells provides a new paradigm that connects membrane biology to the epigenetic regulation of cell plasticity^12^. In particular, increased iron uptake promotes the activity of α-ketoglutarate (αKG)-dependent demethylases involved in the regulation of gene expression. HA have also been shown to induce the expression of pro-inflammatory cytokines in alveolar macrophages^13^, and macrophage activation relies on complex regulatory mechanisms occurring at the chromatin level^14-17^. This body of work raises the question whether a mechanism involving CD44-mediated metal uptake regulates macrophage plasticity.

Here, we show that macrophage activation is characterized by an increase of mitochondrial copper(II), which is mediated by CD44. We discovered that mitochondrial copper(II) catalyzes NAD(H) redox cycling, thereby promoting epigenetic alterations that lead to the inflammatory state. We have developed a small molecule dimer of metformin termed LCC-12 that selectively targets mitochondrial copper(II). This drug induces metabolic and epigenetic shifts that oppose macrophage activation and dampen inflammation *in vivo*.

## Results

### CD44 mediates cellular uptake of Cu

To study the role of metals in immune cell activation, we produced inflammatory macrophages from primary monocytes freshly isolated from blood of human donors (**Fig. 1a**). These cells were differentiated with granulocyte-macrophage colony-stimulating factor (GM-CSF), then treated with lipopolysaccharide (LPS) and interferon gamma (IFNγ) to produce activated monocyte-derived macrophages (aMDM) (**Fig. 1a**). The inflammatory state of aMDM was characterized by the upregulation of the cell surface markers CD80 and CD86, as well as a distinct cell morphology (**Extended Data Fig. 1a-c**). Consistent with the literature^10^, aMDM were also characterized by increased levels of CD44 (**Fig. 1b**).

**Figure 1.**
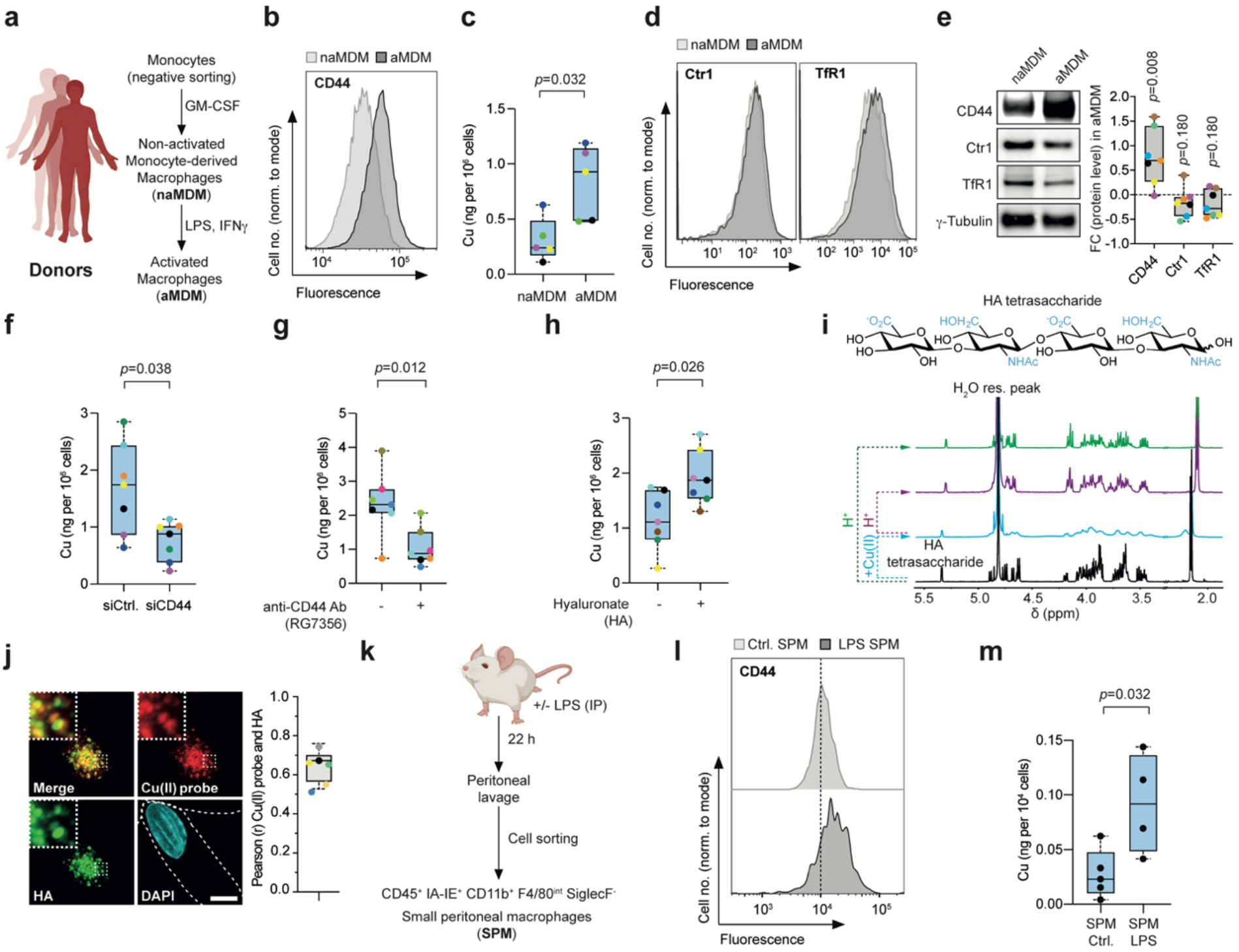
CD44 mediates the uptake of copper. **a**, Experimental setup to generate inflammatory monocyte-derived macrophages (MDM). Peripheral blood samples were collected from 80 donors and were used throughout this study. Pan monocytes were sorted, differentiated with GM-CSF, then activated with LPS and IFNγ to obtain activated MDM (aMDM). Each colored dot represents a specific donor within a given figure panel. **b**, Flow cytometry of CD44 at the plasma membrane. Data representative of *n*=13 donors. **c**, ICP-MS of cellular copper in naMDM and aMDM. *n*=5 donors. **d**, Flow cytometry of Ctr1 and TfR1 at the plasma membrane. Data representative of *n*=3 donors. **e**, Representative western blot of CD44, Ctr1 and TfR1. Quantification displayed on the box-plot. *n*=7 donors. **f**, ICP-MS of cellular copper in aMDM transfected with siCtrl. or siCD44. *n*=7 donors. **g**, ICP-MS of cellular copper in aMDM treated with anti-CD44 antibody RG7356. *n*=7 donors. **h**, ICP-MS of cellular copper in aMDM supplemented with HA (0.6-1 MDa). *n*=7 donors. **i**, Molecular structure of HA tetrasaccharide (top) and ^1^H-NMR spectra (bottom) of copper complexation experiments performed at 37 °C in D_2_O. **j**, Fluorescence microscopy of a lysosomal copper(II) probe and FITC-HA. Dotted lines delineate cell contours. At least 30 cells were quantified per donor. Scale bar, 10 µm. *n*=6 donors. **k**, Scheme of isolation of small peritoneal macrophages (SPM) from the peritoneum of control and LPS-treated mice. A peritoneal lavage was performed after 22 h and SPM were isolated by flow cytometry using the indicated markers. **l**, Flow cytometry of CD44 at the plasma membrane. Data representative of *n*=5 mice per group. **m**, ICP-MS of cellular copper in SPM from control mice and LPS-treated mice. *n*=4-5 mice. Mann-Whitney test for **c, e** – **h**, and **m**. Box plots with median and whiskers of lowest and highest values.

Given that CD44 mediates endocytosis of iron in cancer cells^12^, we investigated whether inflammatory macrophages similarly exploit such a mechanism. Using inductively coupled plasma mass spectrometry (ICP-MS), we observed higher levels of cellular copper, iron, manganese and calcium in aMDM compared to non-activated MDM (naMDM) (**Fig. 1c and Extended Data Fig. 1d**). In contrast, the cellular content of other metals studied, such as magnesium, cobalt, nickel and zinc, was not significantly altered (**Extended Data Fig. 1d**). Interestingly, levels of the copper transporter Ctr1 or the iron transporter TfR1 remained mostly unchanged during macrophage activation (**Fig. 1d, e**). Knocking down CD44 by means of RNA silencing or treating aMDM with the therapeutic anti-CD44 antibody RG7356, which interferes with HA binding to CD44^18^, reduced copper and iron uptake (**Fig. 1f, g and Extended Data Fig. 1e, f**). Conversely, supplementing cells with HA during activation further increased intracellular levels of these metals (**Fig. 1h and Extended Data Fig. 1g**). These data indicate that CD44 and HA promote the uptake of copper and iron in aMDM. To substantiate the role of CD44 in mediating copper uptake, we investigated the propensity of HA to form complexes with copper using a HA tetrasaccharide, the proton signals of which can be resolved by nuclear magnetic resonance (NMR) spectroscopy. Adding copper(II) to HA in water led to a line broadening of the proton signals of HA (**Fig. 1i**). Acidification of the media to protonate the carboxylate of HA and disrupt copper binding restored the signals of unbound HA, showing that the negatively-charged polysaccharides can dynamically interact with copper(II) under physiologically relevant conditions.

Next, we confirmed CD44-dependent copper endocytosis using a fluorescent probe designed to detect copper in lysosomes. Specifically, we used Lys-Cu, whose fluorescence selectively increases upon copper(II) binding^19^. Fluorescence microscopy revealed that a fluorescently labeled HA colocalized with this probe in aMDM (**Fig. 1j**).

Following this, we set out to explore copper levels in a model of inflammation representative of our experimental setup. To this end, we treated mice with LPS intraperitoneally and isolated small peritoneal macrophages (SPM). SPM from mice treated with LPS exhibited higher CD44 levels together with higher copper content compared to healthy mice (**Fig. 1k-m**). Collectively, these data indicate that inflammatory macrophages internalize copper through CD44-mediated HA uptake, raising the hypothesis that copper plays a functional role in macrophage activation.

### Cu(II) regulates cell plasticity

To explore functional roles of copper signaling in the context of inflammation, we evaluated the capacity of the copper chelators D-penicillamine (D-Pen) and ammonium tetrathiomolybdate (ATTM)^20^, to interfere with macrophage activation. In this focused screen we also included metformin (Met), an FDA-approved biguanide used for the treatment of type-2 diabetes, because it forms complexes with copper(II) in cell-free settings^21^. Met partially antagonized CD86 upregulation, albeit at high concentrations (e.g. 10 mM), contrasting the marginal effects of other clinically approved copper-targeting drugs (**Fig. 2a**).

**Figure 2.**
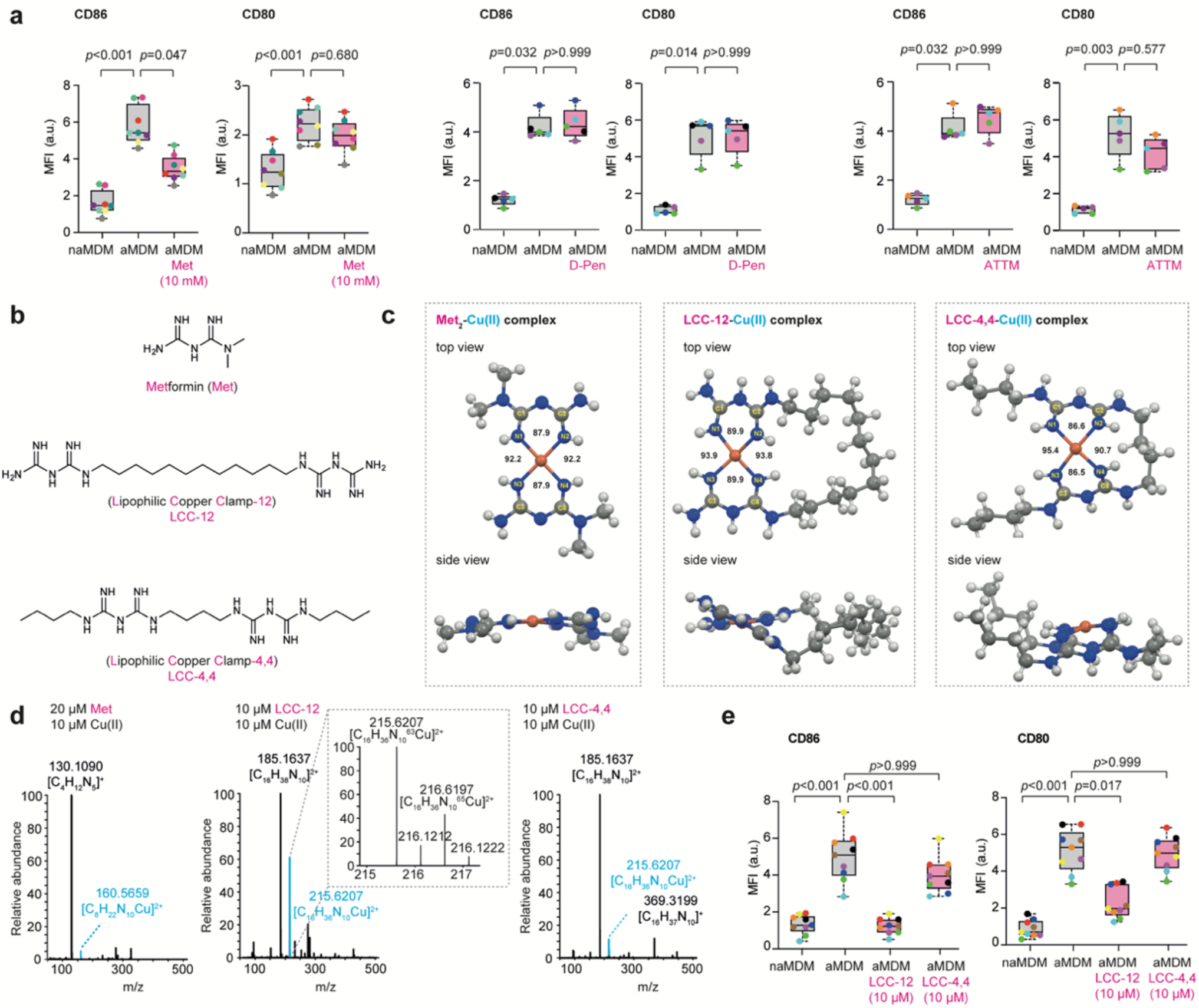
Targeting mitochondrial copper(II) interferes with cell plasticity. **a**, Flow cytometry of CD86 and CD80 cell surface markers of naMDM and aMDM treated with metformin (Met, 10 mM) (*n*=8 donors), D-penicillamine (D-Pen, 250 µM) (*n*=5 donors) or ammonium tetrathiomolybdate (ATTM, 1 µM) (*n*=5 donors). MFI, mean fluorescence intensity. **b**, Molecular structures of Met, LCC-12 and LCC-4,4. **c**, Structural analysis of biguanide-based copper(II) complexes by molecular modeling. Top and side views highlighting the different geometry of Met, LCC-12 and LCC-4,4. **d**, High resolution mass spectrometry (HRMS) of biguanide-based copper(II) complexes. **e**, Flow cytometry of CD86 and CD80 cell surface markers of naMDM, aMDM and aMDM treated with LCC-12 (10 µM) or LCC-4,4 (10 µM). *n*=9 donors. MFI, mean fluorescence intensity. Kruskal-Wallis test with Dunn’s post-test for **a** and **e**. Box plots with median and whiskers of lowest and highest values.

Met can form a stable bimolecular complex with copper(II)^21,22^. To alleviate the entropic cost inherent to the formation of supramolecular complexes and thus, to increase the copper binding capacity of biguanides, we synthesized a series of dimeric small molecules, where two biguanide moieties were tethered with a methylene-containing linker (**Fig. 2b**). The design of such ‘lipophilic copper clamps’ (LCCs) was computationally guided and consisted in varying the number of methylene groups separating two biguanide units to ensure optimal geometry for copper(II) binding. Thus, we synthesized the dimers of biguanides LCC-12 and LCC-4,4 harboring twelve and four methylene-containing linkers, respectively. Importantly, LCC-4,4 was equipped with two distal butyl substituents such that both LCC-12 and LCC-4,4 have identical molecular formulae and exhibit a comparable lipophilicity. We used molecular dynamics simulation to predict the structure of copper(II) complexes with the lowest energies, comparing the geometry of each copper complex with that of an experimentally produced Cu(Met)_2_ complex^22^. This took into account bonding angles around the metal center and the planarity of the system, given that copper(II) complexes can adopt square planar geometries. The modeled geometry of Cu(Met)_2_ resembled that of the crystal structure of a Cu(Met)_2_ complex^22^, validating our theoretical methodology (**Fig. 2c**). Cu-LCC-12 adopted a similar geometry according to bonding angles around copper and the planar geometry of each of the two biguanides bound to the metal ion. These two biguanides occupied distinct planes, whereas Cu(Met)_2_ was planar overall. In contrast, Cu-LCC-4,4 lacked bonding angle symmetry and exhibited imine-copper bonds out of plane, indicating a geometrically constrained structure (**Fig. 2c**). Accordingly, the calculated free energy of Cu-LCC-4,4 was 16.6 kcal/mol higher compared to that of Cu-LCC-12, predictive of a less stable copper(II) complex.

High-resolution mass spectrometry (HRMS) confirmed the formation of monometallic copper complexes with Met, LCC-12 and LCC-4,4 (**Fig. 2d**). Cu-LCC-12 was more stable than Cu(Met)_2_ and Cu-LCC-4,4, as indicated by the height of peaks corresponding to the mass of each complex. Adding copper(II) chloride to a solution of LCC-12 reduced its UV absorbance at 233 nM, indicating complex formation at low micromolar concentrations (**Extended Data Fig. 2a**). Mixing biguanides with copper(II) chloride resulted in colored solutions, which is characteristic of metal complexes (**Extended Data Fig. 2b**). Furthermore, LCC-12 did not form stable complexes with other bivalent metal ions including magnesium, calcium, manganese, zinc and iron (**Extended Data Fig. 2c**). Trace amounts of a LCC-12-Ni complex were detected, and LCC-12 did not form a complex with copper(I). Thus, LCC-12 is a suitable tool for perturbing copper(II) signaling in inflammatory macrophages, while LCC-4,4 can be used as a control analog.

When added to aMDM, LCC-12 antagonized CD80 and CD86 induction more potently than Met, even at a 1000-fold lower dose (e.g. 10 μM LCC-12 versus 10 mM Met) (**Fig. 2e**). In contrast, the effect of LCC-4,4 used at 10 μM was moderate, consistent with the reduced capacity of this analog to interact with copper(II). These data support the idea that copper(II) is a mechanistic target of biguanides.

Next, we evaluated the effect of LCC-12 in other cell types expressing CD44. Interestingly, expression of activation markers by human CD4^+^ or CD8^+^ T lymphocytes, dendritic cells and monocytes, which upregulate CD44 levels upon appropriate stimulation, was reduced upon LCC-12 treatment (**Extended Data Fig. 2d-h**). In contrast, the activation of neutrophils, which did not upregulate CD44, was not affected by LCC-12. Furthermore, in mouse pancreatic adenocarcinoma cells, TGF-β-induced epithelial-mesenchymal plasticity (EMP) was characterized by CD44 upregulation and increased copper levels. In this system, LCC-12 interfered with EMP (**Extended Data Fig. 2i**). Altogether, these data illustrate the general nature of this copper signaling pathway as a regulator of cell plasticity.

### Reactive Cu(II) in mitochondria

To gain further insights into the mechanism of action (MoA) of LCC-12, we employed nanoscale secondary ion mass spectrometry (NanoSIMS) imaging, which qualitatively assesses the cellular distribution of specific isotopes. The subcellular localization of the isotopologue ^15^N-^13^C-LCC-12 overlapped with an ^197^Au-labeled antibody specific for cytochrome *c* (Cyt *c*), suggesting that LCC-12 targets mitochondria in aMDM (**Fig. 3a, b**). To confirm this finding, we developed the biologically active alkyne-containing LCC-12,3 that can be fluorescently labeled in cells by means of click chemistry. This methodology has previously been used to inform on the subcellular localization and putative sites of action of small molecules^23^. In aMDM, the labeled small molecule was detected as cytoplasmic puncta that colocalized with Cyt *c*, confirming accumulation of LCC-12 in mitochondria (**Fig. 3c, d**). The mitochondrial labeling of LCC-12,3 was reduced upon cotreatment with carbonyl cyanide m-chlorophenyl hydrazone (CCCP), a protonophore that dissipates the inner mitochondrial proton gradient (**Fig. 3e**). This indicated that LCC-12 accumulation in mitochondria is driven by its protonation state. Labeling small molecules in cells by means of click chemistry requires a copper(I) catalyst generated *in situ* from added copper(II) and ascorbate (Asc) as a reducing agent. Given that our study converged towards mitochondrial copper(II) as a mechanistic target of LCC-12, we investigated whether the natural abundance of mitochondrial copper in aMDM would allow for click labeling without the need to experimentally add a metal catalyst. We found that fluorescent labeling of LCC-12,3 used at a concentration of 100 nM, which is 100-fold lower than the biologically active dose of LCC-12, occurred in aMDM in absence of exogenous copper, leading to a fluorescent signal that colocalized with mitotracker. This staining pattern was observed only when MDM were activated (**Fig. 3f**), and in the presence of exogenous Asc (**Fig. 3g**). Furthermore, the fluorescence intensity of labeled LCC-12,3 was substantially reduced when a 100-fold molar excess of LCC-12 was used as a non-clickable competitor (**Extended Data Fig. 3a**). ICP-MS measurement using isolated mitochondria revealed that copper levels were higher in aMDM compared to naMDM (**Fig. 3h**). Interestingly, levels of manganese were also increased in mitochondria of aMDM, whereas the contents of other metals studied was not significantly altered (**Extended Data Fig. 3b, c**). Taken together, these data support the existence of a chemically reactive pool of copper(II) in mitochondria of aMDM.

**Figure 3.**
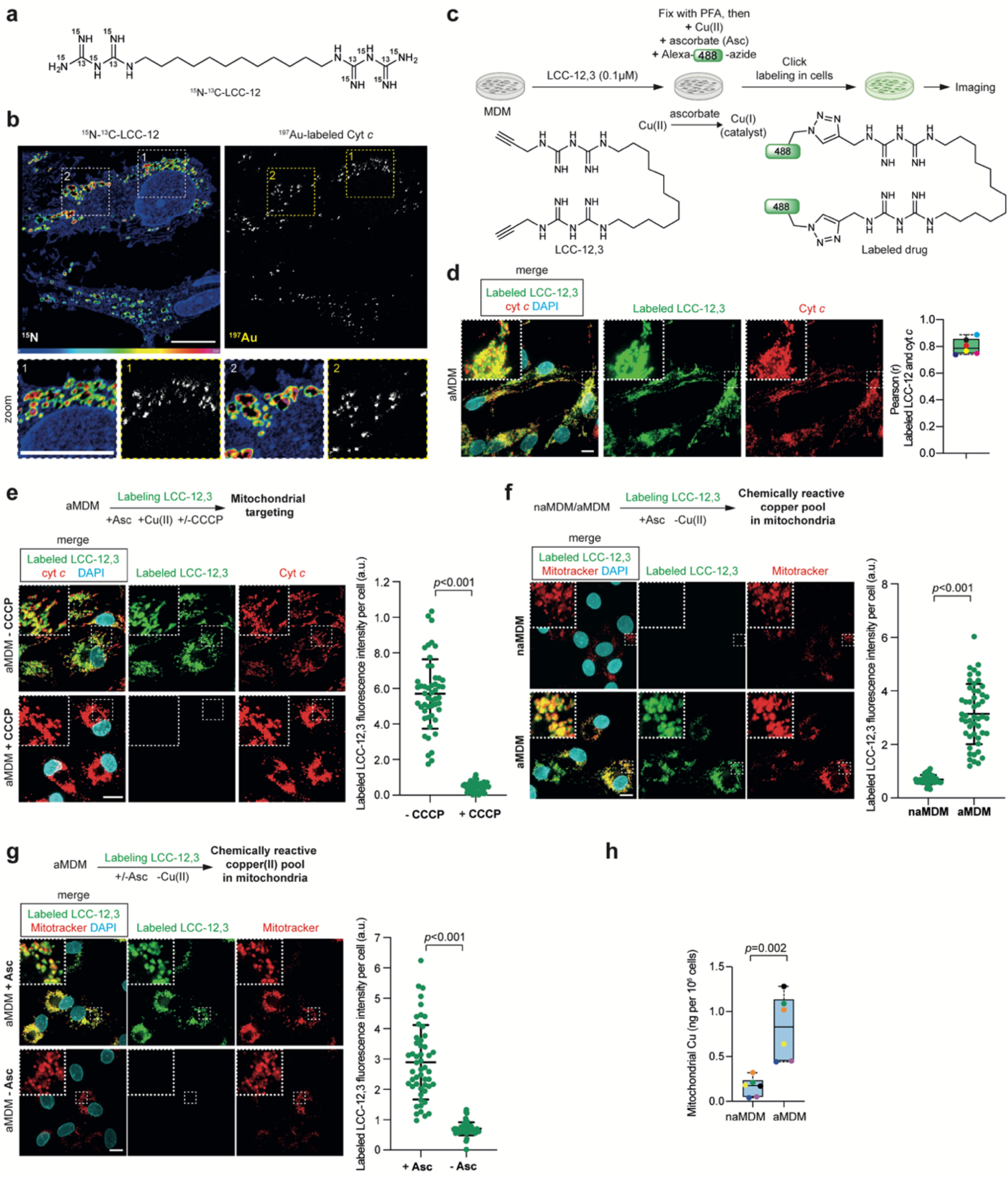
Detection of a druggable pool of copper(II) in mitochondria. **a**, Molecular structure of isotopologue ^15^N-^13^C-LCC-12. **b**, NanoSIMS images of ^15^N and ^197^Au in aMDM. Scale bar, 10 µm. **c**, Schematic illustration of click labeling of alkyne-containing LCC-12,3 in cells. **d**, Fluorescence microscopy of labeled LCC-12,3 (0.1 µM) in aMDM, showing localization in vicinity of cytochrome *c* (Cyt *c*). *n*=6 donors. 50 cells were quantified per donor. **e**, Fluorescence microscopy of labeled LCC-12,3 (0.1 µM) in aMDM cotreated with carbonyl cyanide m-chlorophenyl hydrazone (CCCP). Scale bar, 10 µm. **f**, Fluorescence microcopy images of labeled LCC-12,3 (0.1 µM). Click labeling performed with Asc in absence of added copper(II) in naMDM and aMDM. Scale bar, 10 µm. **g**, Fluorescence microcopy images of labeled LCC-12,3 (0.1 µM). Click labeling performed with or without Asc in absence of added copper(II) in aMDM. Scale bar, 10 µm. **h**, ICP-MS of mitochondrial copper in naMDM and aMDM. *n*=6 donors. Mann-Whitney test. Box plots with median and whiskers of lowest and highest values. **e** – **g** Student’s T-test. Mean ± SEM. At least 50 cells were quantified per condition.

### Cu(II) regulates NAD(H) redox cycling

We next set out to identify mitochondrial processes reliant on copper(II) that contribute to the inflammatory phenotype. In mitochondria, copper is essential for the function of cytochrome *c* oxidase (also termed Complex IV), a component of the electron transport chain (ETC) that catalyzes the reduction of molecular oxygen. Although the regulation of cellular copper homeostasis is not fully understood, it has previously been argued that copper is tightly bound to proteins and mostly handled as a copper(I) species^24^. However, our data suggested that copper(II) plays a critical role in inflammatory macrophages.

Higher mitochondrial levels of manganese in aMDM pointed to a functional role of the superoxide dismutase 2 (SOD2). In line with this, SOD2 protein levels increased in mitochondria upon activation (**Fig. 4a, b**), together with levels of mitochondrial hydrogen peroxide, a product of superoxide dismutation catalyzed by SOD2 (**Fig. 4c, d**). In cell-free systems, copper(II) can catalyze the reduction of hydrogen peroxide using various organic substrates as sources of electrons^25,26^. Mass spectrometry indicated that in the presence of copper(II), hydrogen peroxide reacted with NADH to yield NAD^+^ (**Fig. 4e**). However, in absence of copper, hydrogen peroxide reacted with NADH to yield a complex mixture of products including an epoxide whose structure was supported by spectral data (**Fig. 4e**). Molecular modeling also supported a plausible reaction of epoxidation. In agreement with this, copper(II) favored the conversion of 1-methyl-1,4-dihydronicotinamide (MDHNA), which was used as a structurally less complex surrogate of NADH, into 1-methylnicotinamide (MeNAD^+^), whereas a product of epoxidation formed preferentially in absence of copper (**Extended Data Fig. 4a**). Mass spectrometry revealed the mass of a MDHNA derivative containing an additional oxygen atom, and 2D NMR spectroscopy confirmed epoxidation of the least substituted double bond (**Extended Data Fig. 4b**). Thus, copper reprograms the reactivity of hydrogen peroxide towards NADH, protecting the latter from oxidative degradation. To mimic the conditions found in the mitochondrial matrix, we next performed this reaction at 37 °C and pH 8. The rate of the reaction was evaluated by monitoring the concentration of NADH over time. When the reaction was performed in the presence of copper(II) and imidazole as stabilizing copper ligand, NADH was rapidly consumed to yield NAD^+^ (**Fig. 4f**), as confirmed by mass spectrometry and NMR spectroscopy (**Extended Data Fig. 4c**). This reaction was inhibited by LCC-12 in a dose-dependent manner, whereas the effect of LCC-4,4 and Met was marginal (**Fig. 4f**). Taken together, LCC-12 can effectively compete against hydrogen peroxide for copper(II) binding, thereby inhibiting this copper(II)-catalyzed reaction. Using MDHNA as a model substrate, molecular modeling supported a reaction mechanism involving the formation of a copper(II)-hydrogen peroxide complex, in which copper activates hydrogen peroxide facilitating the transfer of a hydride from NADH onto hydrogen peroxide (**Extended Data Fig. 4d**). Copper(II) can act as a metal catalyst that lowers the energy of the transition state (TS) with a geometry that favors this reaction. Consistent with our findings, molecular modeling also supported that, by forming a complex with copper(II), biguanides can interfere with the binding and activation of hydrogen peroxide (**Extended Data Fig. 4e**).

**Figure 4.**
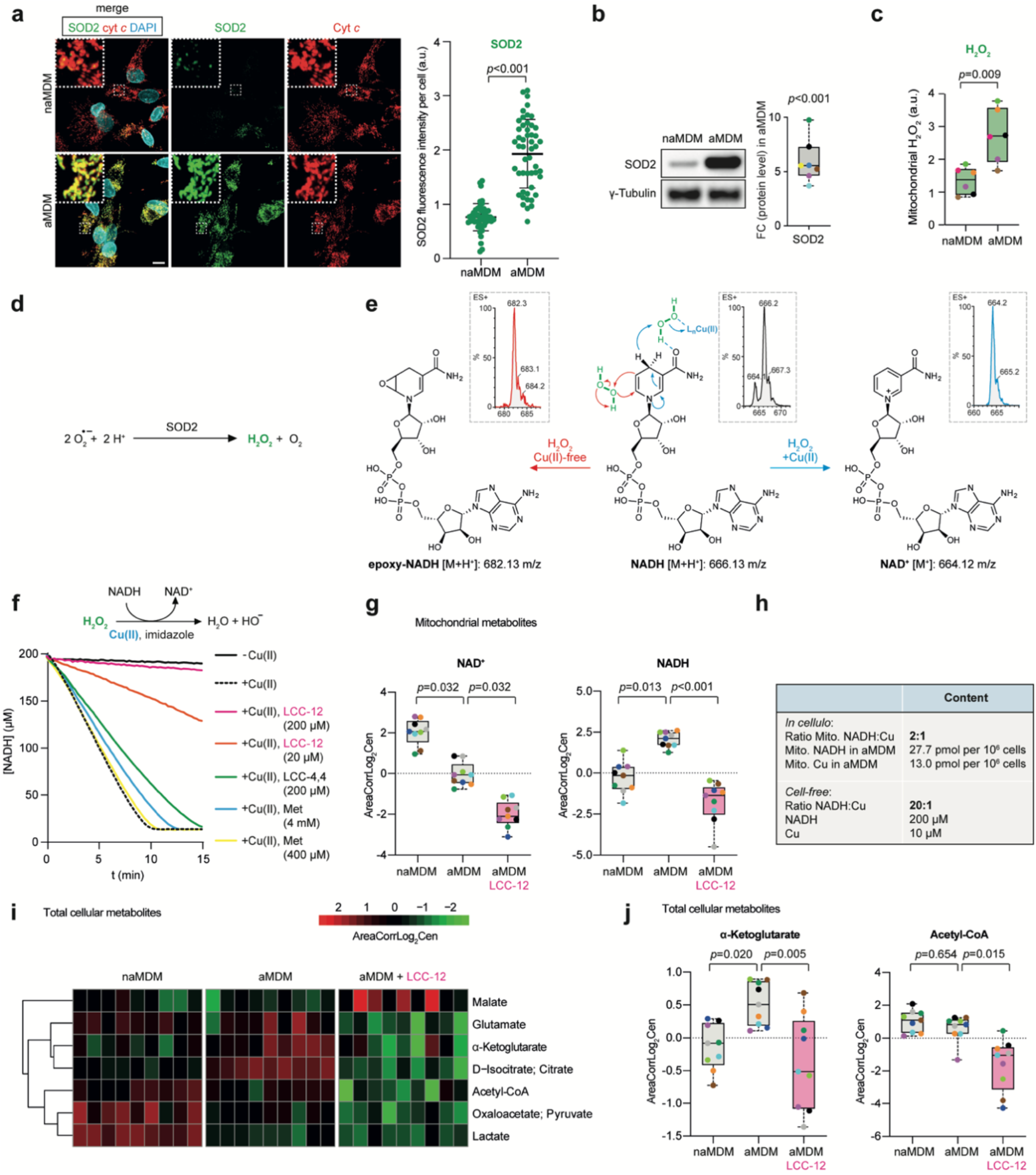
Mitochondrial copper(II) regulates NAD(H) redox cycling. **a**, Fluorescence microcopy images of SOD2 in naMDM and aMDM. Mitochondria were stained using an anti-Cyt *c* antibody. Student’s T-test. Mean ± SEM. At least 50 cells were quantified per condition. **b**, Western blot of SOD2. Data representative of *n*=7 donors. **c**, Flow cytometry analysis of mitochondrial H_2_O_2_ in naMDM and aMDM. *n*=6 donors. **d**, Production of H_2_O_2_ from superoxide catalyzed by superoxide dismutase 2 (SOD2). **e**, Reaction of NADH in the presence of H_2_O_2_ under copper-free and copper(II)-catalyzed conditions. The masses of molecular ions detected by mass spectrometry are indicated. **f**, Copper(II)-catalyzed reduction of hydrogen peroxide by NADH. Kinetics of NADH consumption in the presence of hydrogen peroxide. Data representative of *n*=3 independent replicates. **g**, Quantitative mass-spectrometry-based metabolomics of NAD^+^ and NADH in mitochondrial extracts of naMDM, aMDM and aMDM treated with LCC-12 (10 µM). *n*=9 donors. **h**, Concentrations of copper and NADH in cells and in the cell-free system used in used in **f. i**, Heatmap of quantitative mass-spectrometry-based metabolomics of total cellular extracts highlighting metabolites whose biosynthesis depend on NAD(H). *n*=9 donors. **j**, Box plots of quantitative mass-spectrometry-based metabolomics of αKG and acetyl-CoA. For **a** Student’s T-test. Mean ± SEM. For **b** and **c** Mann-Whitney test. For **g** and **j** Kruskal-Wallis test with Dunn’s post-test. Box plots with median and whiskers of lowest and highest values.

The higher abundance of copper(II) and hydrogen peroxide in mitochondria from aMDM compared to naMDM prompted us to investigate the biological relevance of this reaction in inflammatory macrophages. To this end, we quantified mitochondrial NADH and NAD^+^ in aMDM by mass spectrometry-based metabolomics. Mitochondrial NADH levels were higher, whereas NAD^+^ levels were lower in aMDM compared to naMDM, suggesting an enhanced activity of mitochondrial enzymes reliant on NAD^+^ (**Fig. 4g and Supplementary Table 1**). In agreement with data obtained from our cell-free system, treating aMDM with LCC-12 during activation led to a decrease of both mitochondrial NAD^+^ and NADH (**Fig. 4g and Supplementary Table 1**). This is consistent with the idea that copper(II) catalyzes the reduction of hydrogen peroxide by NADH to produce NAD^+^ in cells, and that biguanides can interfere with this redox cycling, leading instead to other oxidation by-products. Notably, NADH and copper were found in mitochondria of aMDM at an estimated substrate/catalyst ratio of 2:1, which is even more favorable for this reaction to take place than the 20:1 ratio used in the cell-free system (**Fig. 4h**). This substantiates the existence of a copper(II)-catalyzed reduction of hydrogen peroxide by NADH in mitochondria.

Quantitative metabolomics analysis of total cellular extracts indicated that macrophage activation was accompanied by altered levels of several metabolites whose production depend on NAD(H) (**Fig. 4i and Supplementary Table 2**). In particular, LCC-12-induced metabolic reprogramming of aMDM was marked by a reduction of αKG and acetyl-coenzyme A (acetyl-CoA) (**Fig. 4j**). Collectively, these data support the central role of mitochondrial copper(II), which maintains a pool of NAD^+^ and regulates the metabolic state of inflammatory macrophages.

### Cu(II) regulates transcriptional programs

Transcriptional programs underlying macrophage activation involve alterations of the epigenetic landscape^14-17,27^. αKG and acetyl-CoA are key metabolites required to regulate epigenetic plasticity^28^. These two metabolites are co-substrates of iron-dependent demethylases and acetyl transferases (ATs), which can target histones and nucleobases. Our findings that LCC-12 interfered with the production of these metabolites and antagonized macrophage activation, pointed to epigenetic alterations that affect the expression of inflammatory genes. Comparing transcriptional changes triggered by various pathogens can reveal general regulatory mechanisms of macrophage activation. To explore this, we analyzed the transcriptome of aMDM compared to naMDM by RNA-seq (**Supplementary Table 3**), and compared this to transcriptomics data obtained from bronchoalveolar macrophages of patients infected with SARS-CoV-2^29^ and from human macrophages exposed *in vitro* to *Salmonella typhimurium*^30^, *Leishmania major*^31^ or *Aspergillus fumigatus*^32^ (**Supplementary Table 4**). Gene ontology (GO) analysis revealed three main GO term groups, which belonged to inflammation, metabolism and chromatin. Importantly, GO terms of upregulated genes in aMDM included endosomal transport, cellular response to copper ion, response to hydrogen peroxide and positive regulation of mitochondrion organization (**Fig. 5a**). Furthermore, we observed striking similarities between the transcriptomics datasets (**Extended Data Fig. 5a, b and Supplementary Table 5**). In particular, we found increased RNA levels of genes encoding inflammatory cytokines such as IL-6, IL-1β and TNFα, as well as genes encoding proteins involved in the JAK/STAT signaling pathway, the inflammasome and Toll-Like Receptors (TLRs) (**Fig. 5b and Extended Data Fig. 5c**). In addition, aMDM upregulated genes that encode sorting nexin 9 (SNX9), a regulator of CD44 endocytosis^33^, the lysosomal copper transporter 2 (CTR2) and metallothioneins (MT2A, MT1X), which are involved in copper transport and storage (**Supplementary Table 3**). In contrast, the gene coding for COX11, a chaperone involved in copper incorporation into cytochrome *c* oxidase was downregulated.

**Figure 5.**
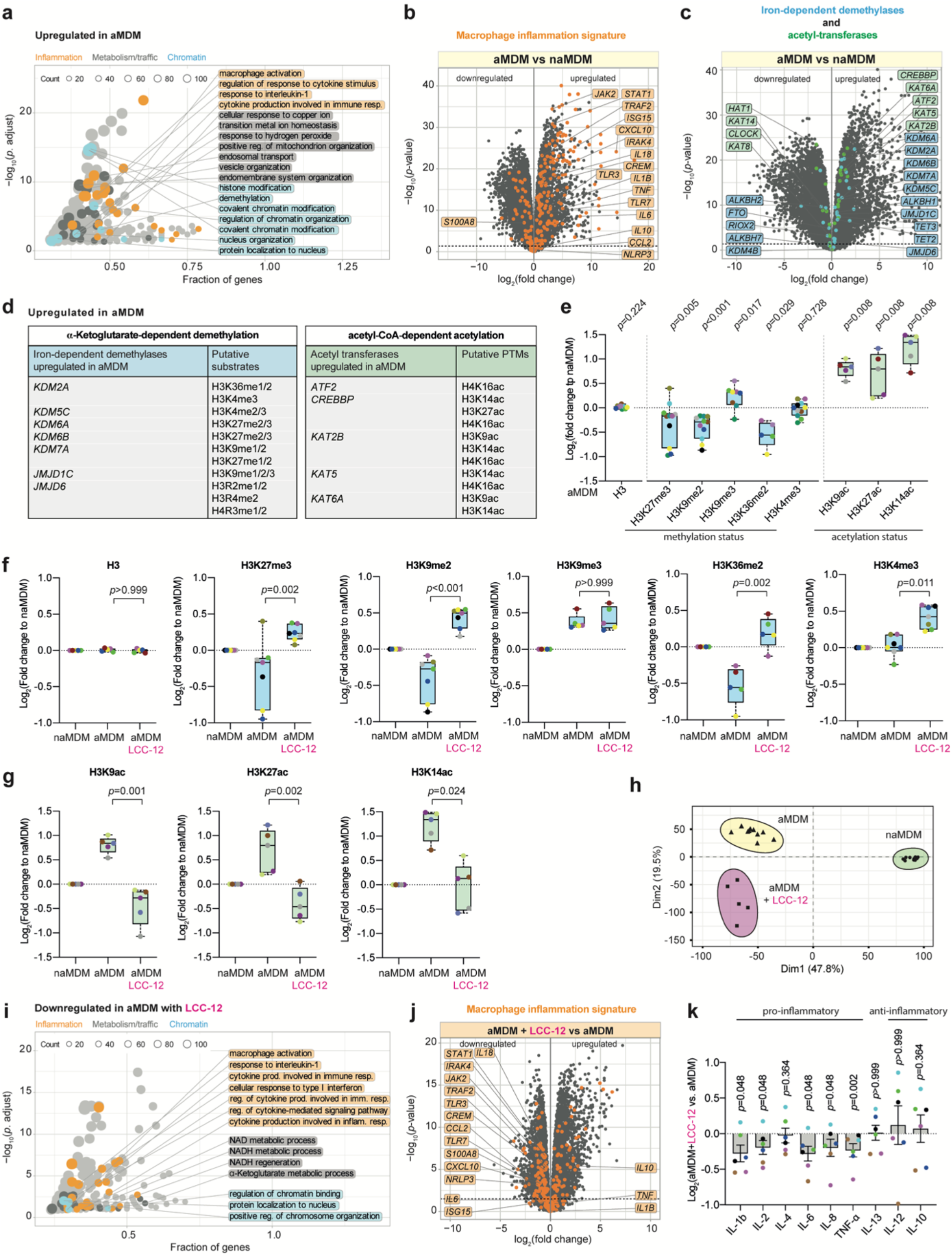
Mitochondrial copper(II) regulates the epigenetic state and transcriptional programs. **a**, GO term analysis of upregulated genes in aMDM versus naMDM. **b**, RNA-seq analysis of gene expression in aMDM versus naMDM. Inflammatory gene signature is illustrated (orange). Dashed line, adjusted *p*-value=0.05. **c**, RNA-seq analysis of gene expression in aMDM versus naMDM. Genes encoding iron-dependent demethylases (blue) and histone acetyl-transferases (green) are illustrated. Dashed line, adjusted *p*-value=0.05. **d**, Genes coding for iron-dependent demethylases and acetyl-transferases found to be upregulated in aMDM are listed. Putative substrates are listed. **e**, Quantifications of histone H3 methyl and acetyl marks regulated by the enzymes overexpressed in aMDM using fluorescence microscopy. Quantifications represent aMDM normalized against naMDM. At least 50 cells were quantified per donor per condition. *n*=5-11 donors. Mann-Whitney test. Box plots with median and whiskers of lowest and highest values. **f**, Quantifications of histone H3 methyl marks in aMDM treated with LCC-12 (10 µM) using fluorescence microscopy. Quantifications represents aMDM normalized against naMDM. At least 50 cells were quantified per donor per condition. *n*=5-7 donors. Kruskal-Wallis test with Dunn’s post-test. Box plots with median and whiskers of lowest and highest values. **g**, Quantifications of histone H3 acetyl marks in aMDM treated with LCC-12 (10 µM) using fluorescence microscopy. Quantifications represent aMDM normalized against naMDM. At least 50 cells were quantified per donor per condition. *n*=5 donors. Kruskal-Wallis test with Dunn’s post-test. Box plots with median and whiskers of lowest and highest values. **h**, Principal Component Analysis (PCA) of RNA-seq comparing naMDM (*n*=10 donors), aMDM (*n*=10 donors) and aMDM treated with LCC-12 (10 µM) (*n*=5 donors). **i**, GO term analysis of genes in aMDM whose expression is downregulated upon treatment with LCC-12. **j**, RNA-seq analysis of gene expression in aMDM treated with LCC-12 (10 µM) versus aMDM. Inflammatory gene signature is illustrated (orange). Dashed line, adjusted *p*-value=0.05. **k**, Luminex immunoassay of cytokines in the supernatant of aMDM and aMDM treated with LCC-12 (10 µM) (*n*=6 donors). Mann-Whitney test. Error bars represent SEM.

Next, we explored epigenetic gene signatures and found that genes involved in chromatin and histone modifications were upregulated in aMDM (**Fig. 5a, c, d**). A similar subset of genes encoding iron-dependent demethylases and ATs was upregulated in bronchoalveolar macrophages of patients infected by SARS-CoV-2 and in macrophages exposed to other pathogens (**Extended Data Fig. 5d**). These data indicate that macrophage activation relies on similar epigenetic reprogramming irrespective of the pathogens involved^34^ and is a key element of macrophage activation. In aMDM, we observed a reduction of H3K9me2, H3K27me3 and H3K36me2, which are substrates of iron-dependent demethylases (**Fig. 5e**). We also found an increase of H3K9ac, H3K14ac and H3K27ac, which are products of ATs. It is noteworthy that the higher iron load detected in aMDM is consistent with an increased demethylase activity, which contributes to establishing the inflammatory phenotype. Depletion of repressive marks (e.g. H3K9me2, H3K27me3) and increase of marks associated with active transcription (e.g. H3K9ac, H3K14ac, H3K27ac) provide a mechanistic rationale underlying the expression of inflammatory genes in macrophages.

LCC-12 treatment led to an increase of methylation and a reduction of acetylation of specific histone residues including H3K9 and H3K27 (**Fig. 5f, g**). Thus, LCC-12-induced decrease of αKG and acetyl-CoA reduced the activity of iron-dependent demethylases and ATs, respectively. Accordingly, LCC-12 altered the gene expression signature of aMDM (**Supplementary Table 3**), reflecting a complex epigenetic reprogramming toward a distinct cell state (**Fig. 5h**). Notably, targeting mitochondrial copper(II) downregulated genes related to NAD(H) and αKG metabolism, regulation of chromatin and inflammation (**Fig. 5i and Supplementary Table 6**). In aMDM, LCC-12 induced a downregulation of genes encoding IL6, STAT1, JAK2, components of the inflammasome and TLRs among others (**Fig. 5j**). Consistently, secretion of pro-inflammatory cytokines was reduced (**Fig. 5k**). Altogether, these data advocate for a biological mechanism whereby inflammatory macrophages increase mitochondrial copper(II) levels to replenish the pool of NAD^+^. This process is involved in the production of αKG and acetyl-CoA, metabolites that are required for the epigenetic regulation of inflammatory gene expression.

### Cu(II) inactivation reduces inflammation

Next, we investigated the biological effect of LCC-12 in mouse models of bacterial infection, where inflammatory macrophages play a central role^2,7^. At a dose of 3 mg/kg by intraperitoneal (IP) injection, LCC-12 did not provoke any adverse clinical signs. In contrast, at 10 mg/kg, signs of piloerection that vanished overtime were observed in 60% of the cohort, whereas additional signs of ptosis, reduced activity and death occurred in 40% of the cohort. We measured a maximal bioavailability of 61% fifteen minutes after intraperitoneal injection. LCC-12 was stable in plasma and exhibited moderate binding to plasma proteins.

First, we evaluated the effect of LCC-12 at 0.3 mg/kg/day, a dose at least 10 times lower than the maximum tolerated dose (MTD), in two well-established models of sepsis, namely (i) endotoxemia induced by LPS and (ii) cecal ligation and puncture (CLP). The former was chosen to reflect our mechanistic model of macrophage activation, whereas the latter is representative of the pathophysiology of subacute polymicrobial abdominal sepsis occurring in humans^35^. Like SPM from mice treated with LPS (**Fig. 1m**), peritoneal tissues exhibited higher levels of copper after local injection of LPS (**Fig. 6a**). Treatment with LCC-12 fully protected animals from LPS-induced death and prevented the reduction of body temperature (**Fig. 6b, c**). It is noteworthy that LCC-12 performed better than high doses of the anti-inflammatory glucocorticoid dexamethasone (DEX), which is used for the clinical management of acute inflammation and severe forms of COVID-19^5^. Importantly, LCC-12 treatment decreased the inflammatory state of SPM *in vivo* (**Fig. 6d**). Consistently, in a model of CLP-induced lethal sepsis, LCC-12 delayed death and increased the survival rate of mice, comparing favorably to DEX (**Fig. 6e**).

**Figure 6.**
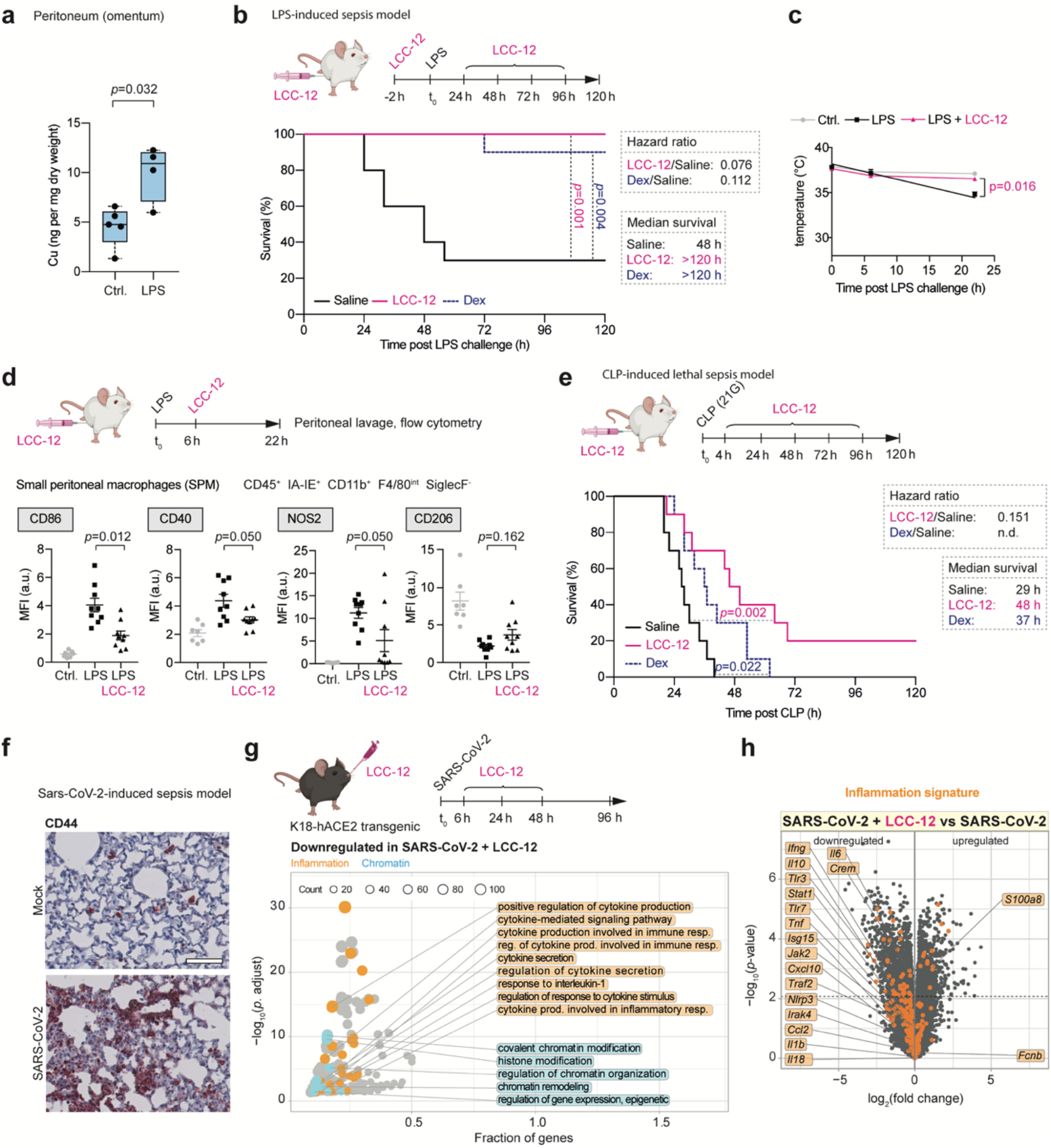
Pharmacological inactivation of copper attenuates inflammation in mouse models of bacterial and viral infection. **a**, ICP-MS of copper of omenta from mice challenged with LPS. *n*=4-5. Mann-Whitney test. Box plots with median and whiskers of lowest and highest values. **b**, Kaplan-Meier survival curves of mice challenged with LPS (20 mg/kg/single dose, IP, *n*=10) and treated with LCC-12 (0.3 mg/kg, 2 h prior challenge, then 24 h, 48 h, 72 h and 96 h post challenge, IP, *n*=10) or dexamethasone (10 mg/kg/single dose 1 h prior challenge, *per os* (PO), *n*=10). Mantel-Cox log-rank test. Hazard ratio calculated using Mantel-Haenszel. **c**, Average temperature in LPS-induced sepsis model. *n*=6-9 mice per group. 2-way ANOVA. Mean values ± SEM. **d**, Flow cytometry of intracellular NOS2 and cell surface markers in SPM from mice challenged with LPS and treated with LCC-12. *n*=7-9 mice per group. Kruskal-Wallis test with Dunn’s post-test. Mean ± SEM. MFI, mean fluorescence intensity. **e**, Kaplan-Meier survival curves of mice subjected to CLP using a 21G needle and treated with LCC-12 (0.3 mg/kg, 4 h, 24 h, 48 h, 72 h and 96 h post-CLP, IP, *n*=10), dexamethasone (1.0 mg/kg at t_0_) or saline solution (IP, *n*=10). Mantel-Cox log-rank test. Hazard ratio calculated using Mantel-Haenszel. n.d. = not determined. **f**, CD44 immunohistochemical staining of lungs from mice infected with SARS-CoV-2. Scale bar = 100 µm. **g**, GO term analysis of genes whose expression is downregulated upon treatment with LCC-12 in SARS-Cov-2 infected mice. **h**, RNA-seq analysis of gene expression in lung tissues of SARS-CoV-2-infected mice treated with LCC-12 versus untreated. Inflammatory gene signature is illustrated (orange). Dashed lines, adjusted *p*-value=0.05.

Next, we evaluated the effect of LCC-12 in a model of viral infection. K18-hACE2 transgenic mice infected with SARS-CoV-2 displayed increased levels of CD44 in lung tissues (**Fig. 6f**). Similarly, lung tissues were characterized by alterations of genes related to mitochondrial metabolism, regulation of chromatin and inflammation (**Extended Data Fig. 6a and Supplementary Table 7**). In particular, viral infection promoted the expression of inflammatory genes and some genes encoding ATs and iron-dependent demethylases (**Extended Data Fig. 6b, c and Supplementary Table 8**). Treatment with LCC-12 administered by inhalation perturbed the expression of genes involved in the regulation of chromatin and inflammation (**Fig. 6g and Supplementary Table 9**), with a reduced expression of inflammatory genes (**Fig. 6h, Extended Data Fig. 6d and Supplementary Table 8**). Taken together, these data indicate that targeting mitochondrial copper(II) interferes with acquisition of the inflammatory state *in vivo* and confers therapeutic benefits.

## Discussion

We have uncovered a copper-signaling pathway that regulates the plasticity of macrophages toward an inflammatory state. We report that the pro-inflammatory cell surface marker CD44 mediates copper uptake to promote macrophage activation. We provide evidence for the existence of a labile pool of copper(II) in mitochondria that characterizes the inflammatory cell state and describe a previously uncharted chemical reaction that takes place in this organelle. We found that NAD^+^ maintenance enables epigenetic programming to unlock the expression of inflammatory genes. Transcriptomics data of macrophages exposed to distinct classes of pathogens, including bacteria and viruses, illustrate the general nature of this mechanism. Macrophages play multiple roles in health and disease. For instance, inflammation is part of the etiology of a plethora of diseases and biological processes such as cancer, aging and obesity^36^. Thus, unraveling and manipulating mechanisms underpinning cell activation and plasticity is crucial for the design of novel therapeutic strategies. We show that mitochondrial copper(II) is druggable using a rationally-designed small molecule, which leads to increased survival and improved well-being in murine models of sepsis.

The cell surface marker CD44 has previously been linked to inflammation, immune response and cancer progression. Here, we show that CD44 mediates cellular uptake of copper, which regulates cell plasticity in distinct cell types. Thus, CD44 can be more generally defined as a regulator of cell plasticity. Copper has previously been shown to play a role in immune defense and cancer^37,38^. For example, copper has been proposed to promote the production of reactive oxygen species as a defense mechanism against bacteria^39^. Furthermore, targeting copper trafficking was reported to attenuate cancer cell proliferation^40^.

Our data indicate that the mitochondrial copper(II) content increases during macrophage activation and that this pool can be targeted with small molecules. Furthermore, inflammatory macrophages exhibit increased levels of hydrogen peroxide in mitochondria. While this substrate is often described as a potentially toxic by-product of respiration, our data suggest a distinct role. Macrophages upregulate the production of hydrogen peroxide to replenish the pool of NAD^+^ enzymatic cofactor from NADH, which may be energetically more advantageous for cells compared to *de novo* biosynthesis. In this context, hydrogen peroxide is a functional metabolite. Our data support the existence of a chemical reaction in which copper(II) activates hydrogen peroxide in mitochondria, enabling its reduction by NADH through a hydride transfer to yield NAD^+^. Conversely, in absence of copper, the reaction of NADH and hydrogen peroxide yields by-products of oxidation. Thus, in inflammatory macrophages, copper(II) protects NADH from degradation, while maintaining NAD(H) redox cycling. This is consistent with the reactivity of enamines (e.g. NADH, MDHNA) towards dimethyldioxirane, an organic equivalent of hydrogen peroxide that can promote the production of highly reactive epoxides prone to solvolysis and alkylation^41^. Interestingly, epoxide hydrolases have been detected in the mitochondria of mammalian cells to prevent accumulation of these chemically reactive species that can potentially lead to alkylation by-products^42^.

Manganese is exploited by SOD2 for the dismutation of superoxide^43^, and iron serves as a metal catalyst of demethylases consuming molecular oxygen^44^. This raises the question of whether this copper-catalyzed reaction is similarly assisted by an enzyme. The experimental setup we employed indicates that this reaction can be performed in an enzyme-free manner. Imidazole, a functional group of histidine (His) residues found in mitochondrial proteins, enhances the rate of this reaction through binding to copper(II), arguing in favor of a putative enzyme-mediated process. A plausible hypothesis points to complex I of the ETC, which contains a NADH binding site and imidazole-containing His residues in proximity of flavin mononucleotide (FMN)^45^, being involved in the reduction of hydrogen peroxide by NADH. There, copper(II) bound to His may activate hydrogen peroxide with a geometry poised for hydride transfer. This is further supported by the observation that Met and a copper(II)-thiosemicarbazone complex inhibit complex I of the ETC^46,47^. Complex I may accommodate other substrates than FMN, promoting reduction of hydrogen peroxide.

Metals are ubiquitous in cell biology, serving multiple functions. While other processes may be at play, our data illuminate a central role of copper in the activation of macrophages by promoting the expression of inflammatory genes. This involves NAD^+^-dependent production of αKG and acetyl-CoA, two metabolites required for the regulation of epigenetic plasticity. These metabolites enable demethylation and acetylation of chromatin marks to unlock the expression of inflammatory genes. We identified a series of iron-dependent demethylases and acetyl transferases reliant on these co-substrates that were upregulated in aMDM and macrophages exposed to various pathogens, suggesting a common epigenetic mechanism underlying macrophage activation.

We designed a dimer of biguanides able to form a near-square planar copper(II) complex. Upon binding to copper(II), LCC-12 prevents activation of hydrogen peroxide, which reduces levels of NAD(H) and interferes with αKG and acetyl-CoA biosynthesis. This in turn impairs the enzymatic activity of specific demethylases and ATs, causing increased histone methylation and a reduced expression of inflammatory genes. LCC-12 exhibits therapeutic effects in several models of acute inflammation, illustrating the pathophysiological relevance of this copper(II)-triggered molecular chain of events. Acute inflammation is therefore reminiscent of a metabolic disease that can be rebalanced by controlling cell plasticity through the targeting of mitochondrial copper(II) (**Extended Data Fig. 7**).

A remarkable feature of LCC-12 is to selectively target mitochondrial copper(II), a unique property that may not be achieved by genetic manipulation or other clinically approved small molecules. Regardless, it appears that the metformin-inspired mitochondrial copper(II) interactor LCC-12 may serve as a lead compound for developing anti-inflammatory agents based on this MoA. Due to its broad anti-inflammatory action, metformin exhibits positive effects on human health and is actually being studied as an anti-aging drug in a large NIH-sponsored human clinical trial^48^. However, exploration of the MoA of metformin is hampered by its poor pharmacology resulting in low potency, which impose high administration doses *in vitro* and *in vivo*^49,50^. Hence, more potent biguanide derivatives such as LCC-12 provide the means to characterize and eventually improve on the clinical and biological characteristics of this class of drugs. Overall, our findings illuminate the central role of copper as a regulator of cell plasticity and unveil a new therapeutic strategy based on the control and fine-tuning of epigenetic cell states.

## Acknowledgements

RR expresses his gratitude to J.-M. Lehn whose inspiring achievements has been instrumental to this work. This work was supported in part by INSERM and CNRS. We thank the PICT-IbiSA@BDD Imaging Facility of Institut Curie, member of the France-BioImaging national research infrastructure (ANR-10-INBS-04) for the use of microscopes, the cytometry platform of Institut Curie, C. Gaillet, J. Sampaio Lopes and D. Guillemot for technical assistance, the ICP-MS platform at Institut de Physique du Globe de Paris, K. Bailly from the Cytometry and Immuno-Biology (CYBIO) platform of Institut Cochin. G.D.P. thank the University of Bath for support. This research made use of the Balena High Performance Computing (HPC) Service at the University of Bath. V.G. thanks the UPSaclay, CNRS and Ecole Polytechnique for financial support. This work has been funded by the following: The European Research Council under the European Union’s Horizon 2020 research and innovation programme grant agreement No 647973 (RR), The Foundation Charles Defforey-Institut de France (RR), Ligue Contre le Cancer (RR and GK Equipes Labellisées), Region IdF for NMR infrastructure and financial support (RR), FHU Sepsis and RHU records (DA), Agence National de la Recherche Projets blancs AMMICa US23/CNRS UMS3655, Association pour la recherche sur le cancer, Association “Ruban Rose”, Cancéropôle Ile-de-France, Fondation pour la Recherche Médicale, A donation by Elior, Equipex Onco-Pheno-Screen, European Joint Programme on Rare Diseases (EJPRD), Gustave Roussy Odyssea, The European Union Horizon 2020 Projects Oncobiome and Crimson, Fondation Carrefour, Institut National du Cancer (INCa), Institut Universitaire de France, LabEx Immuno-Oncology (ANR-18-IDEX-0001), The Leducq Foundation, The RHU Torino Lumière, Seerave Foundation, SIRIC Stratified Oncology Cell DNA, Repair and Tumor Immune Elimination (SOCRATE), SIRIC Cancer Research and Personalized Medicine (CARPEM), This study contributes to the IdEx Université de Paris ANR-18-IDEX-0001, Institut de Physique du Globe de Paris is supported by IPGP multidisciplinary program PARI and Paris–Region IdF (SESAME grant agreement No [12015908]), This work was granted access to the HPC resources of CINES under the allocation 2020-A0070810977 made by GENCI, FT thanks for the support by University of Lille and Institut Pasteur de Lille.

## Author contributions

R.R. conceptualized the study and directed the research. R.R., S.M., S.S. and T.C. designed the experiments and analyzed the data. S.S. performed monocyte isolation, flow cytometry, metabolomics, RNA-seq and cytokine titration. S.M. performed western blotting, brightfield and fluorescence imaging, and ICP-MS. T.C., A.V. and L.B. synthesized the small molecules and fluorescent probes, performed NMR spectroscopy, high-resolution mass spectrometry, UV measurements, and conducted copper-catalyzed NADH oxidation kinetics. G.D.P. and V.G. performed molecular modeling. T.-D.W. performed NanoSIMS imaging. P.G. and N.S. processed RNA-seq data. L.E., A.M., V.S., C.R., F.T. and D.A. conducted *in vivo* experiments and PCR analysis. A.-L.B. and H.S. performed immunohistochemistry. S.D. assisted with quantitative metabolomics. R.R., S.M. and S.S. wrote the article. N.M., A.P., S.W., M.A.D., G.K., and D.A. commented on the manuscript.

## Data and code availability

RNA-seq data are available on the National Center for Biotechnology Information website with accession reference GSE160864 (https://www.ncbi.nlm.nih.gov/geo/query/acc.cgi?acc=GSE160864; enter token wvqxgwgojxcdboz into the box). Analysis scripts are available at https://github.com/bioinfo-pf-curie/MDMmetals All data are available in the main text or the supplemental materials.

## Supplementary information

Extended Data Figures 1-7

Supplementary Tables 1-9

Materials and Methods

Supplementary references 51-78

