## Supplementary Information for "Discovery of a druggable copper-signaling pathway that drives cell plasticity and inflammation"

**This document includes:**

Extended Data Figures 1-7
Captions for Supplementary Tables 1-9
Materials and Methods
Supplementary References

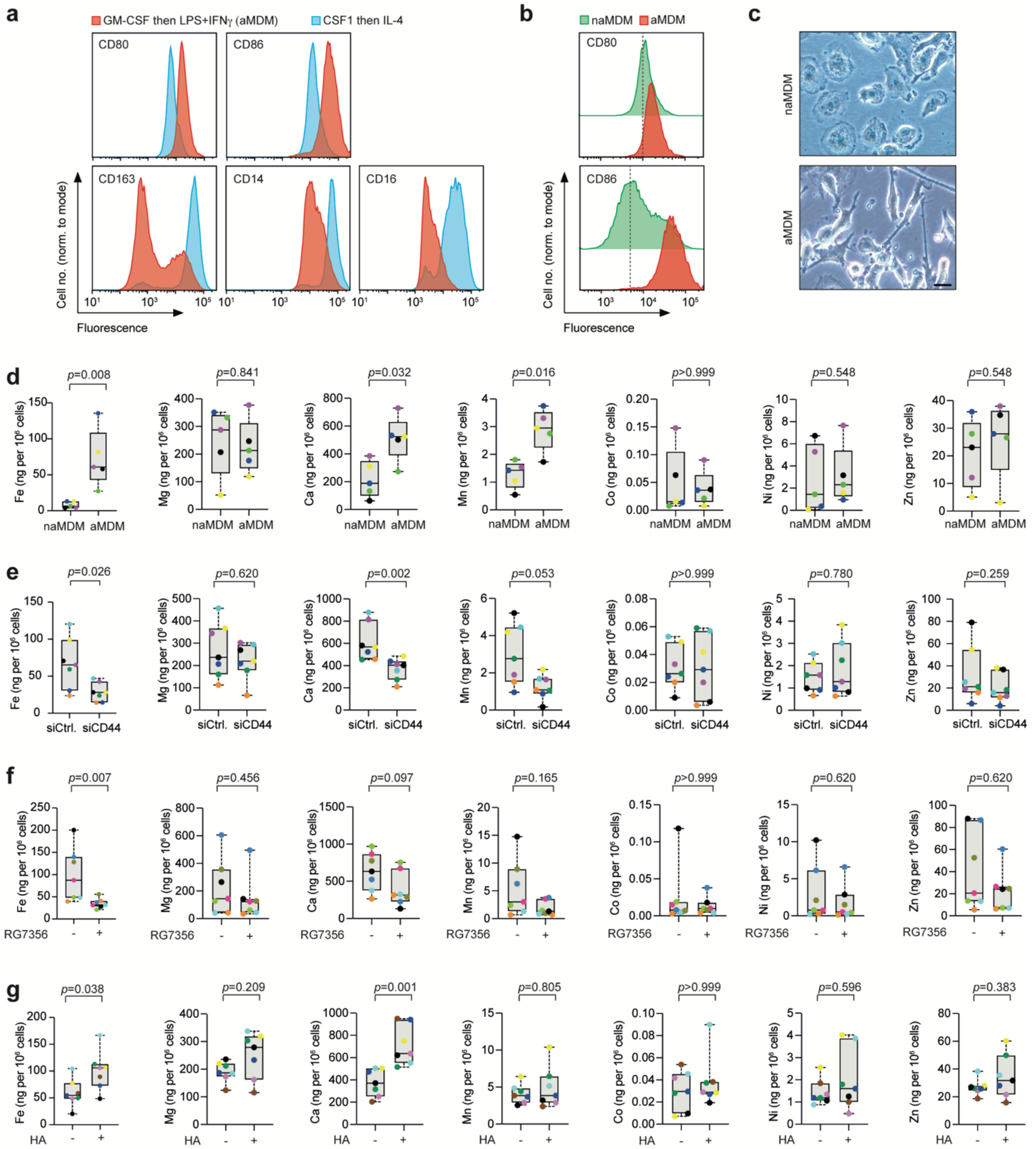


**Extended Data Figure 1 | CD44-mediated iron uptake in inflammatory macrophages. a**, Flow cytometry of cell surface markers of macrophages. Monocytes were differentiated with GM-CSF, then activated with LPS and IFNγ to obtain inflammatory macrophages. For comparison, monocytes were differentiated with CSF1 and then activated with IL-4 to obtain differentially polarized macrophages. **b**, Flow cytometry of CD80 and CD86 cell surface markers of naMDM and aMDM. Data representative of $n$=13 donors. **c**, Bright field microscopy images of aMDM. Scale bar, 20 µm. **d**, ICP-MS of cellular metals in naMDM and aMDM. $n$=5 donors. **e**, ICP-MS of cellular metals in aMDM transfected with siCtrl. or siCD44. $n$=7 donors. **f**, ICP-MS of cellular metals in aMDM treated with an anti-CD44 antibody. $n$=7 donors. **g**, ICP-MS of cellular metals in aMDM supplemented with HA (0.6-1 MDa). $n$=7 donors. For **d** – **g** Mann-Whitney test. Box plots with median and whiskers of lowest and highest values.

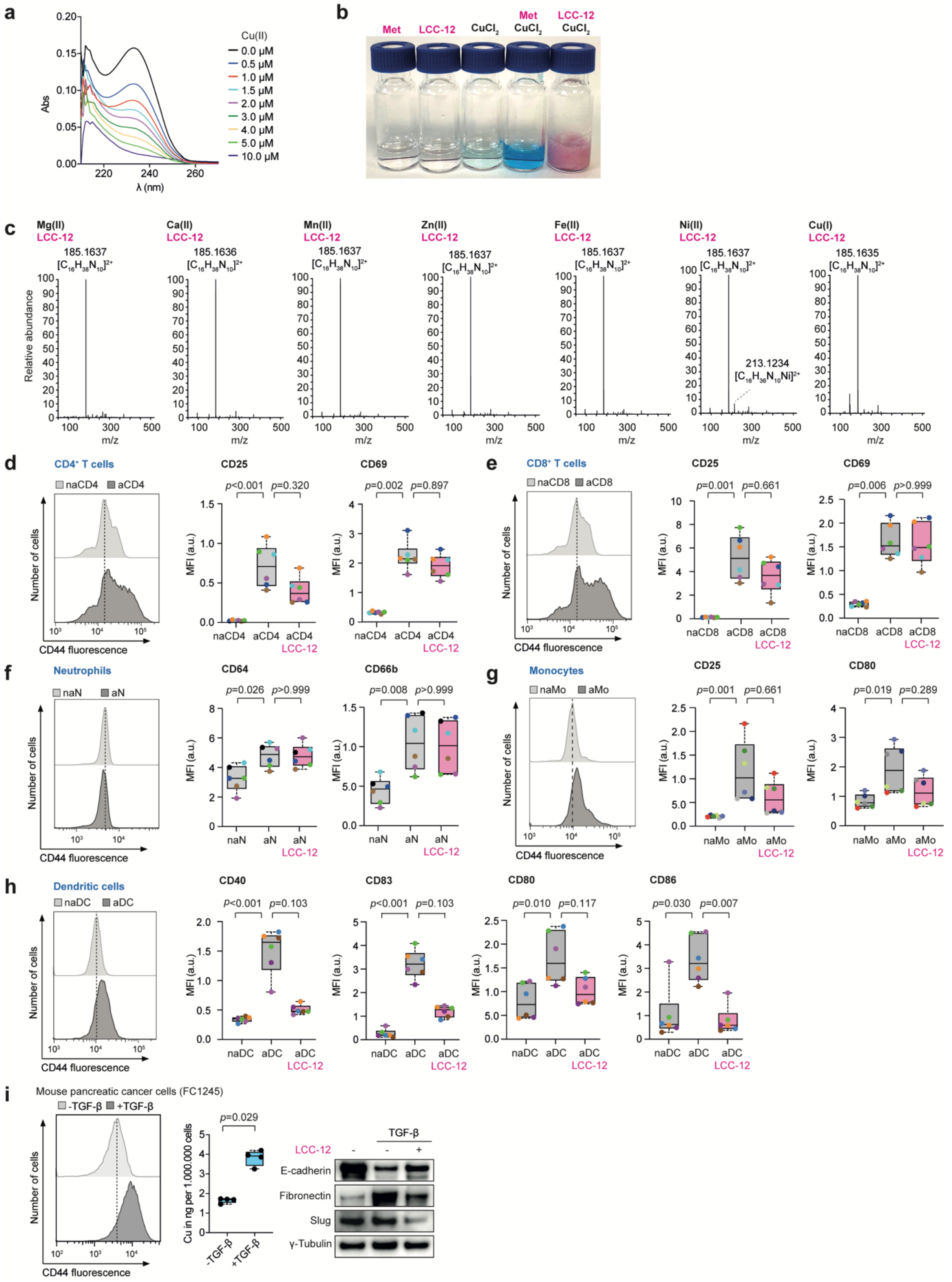


**Extended Data Figure 2 | Targeting mitochondrial copper(II) interferes with cell plasticity. a**, UV spectra of LCC-12 (5 µM in HEPES buffer) titrated with a solution of copper(II). **b**, Picture of aqueous solutions of Met, LCC-12, $CuCl_2$ and corresponding mixtures. **c**, High resolution mass spectrometry (HRMS) of LCC-12 in the presence of metals as indicated. **d**, $CD4^+$ T cells, non-activated (naCD4) and activated (aCD4). Left: Flow cytometry of CD44 at the plasma membrane. Right: Flow cytometry of CD25 and CD69 cell surface markers of naCD4, aCD4 and aCD4 treated with LCC-12 (10 µM). **e**, $CD8^+$ T cells, non-activated (naCD8) and activated (aCD8). Left: Flow cytometry of CD44 at the plasma membrane. Right: Flow cytometry of CD25 and CD69 cell surface markers of naCD8, aCD8 and aCD8 treated with LCC-12 (10 µM). **f**, Neutrophils, non-activated (naN) and activated (aN). Left: Flow cytometry of CD44 at the plasma membrane. Right: Flow cytometry of CD64 and CD66b cell surface markers of naN, aN and aN treated with LCC-12 (10 µM). **g**, Monocytes, non-activated (naMo) and activated (aMo). Left: Flow cytometry of CD44 at the plasma membrane. Right: Flow cytometry of CD25 and CD80 cell surface markers of naMo, aMo and aMo treated with LCC-12 (10 µM). **h**, Dendritic cells, non-activated (naDC) and activated (aDC). Left: Flow cytometry of CD44 at the plasma membrane. Right: Flow cytometry of CD40, CD83, CD80 and CD86 cell surface markers of naDC, aDC and aDC treated with LCC-12 (10 µM). **i**, Mouse pancreatic cancer cells (FC1245) +/- TGF-β. Left: Flow cytometry of CD44 at the plasma membrane. Right: Western blot of FC1245 cells treated with TGF-β and LCC-12 (1 µM, 3d) as indicated. For **d** – **h** Kruskal-Wallis test with Dunn's post-test. For **i** Mann-Whitney test. Box plots with median and whiskers of lowest and highest values. MFI, mean fluorescence intensity.

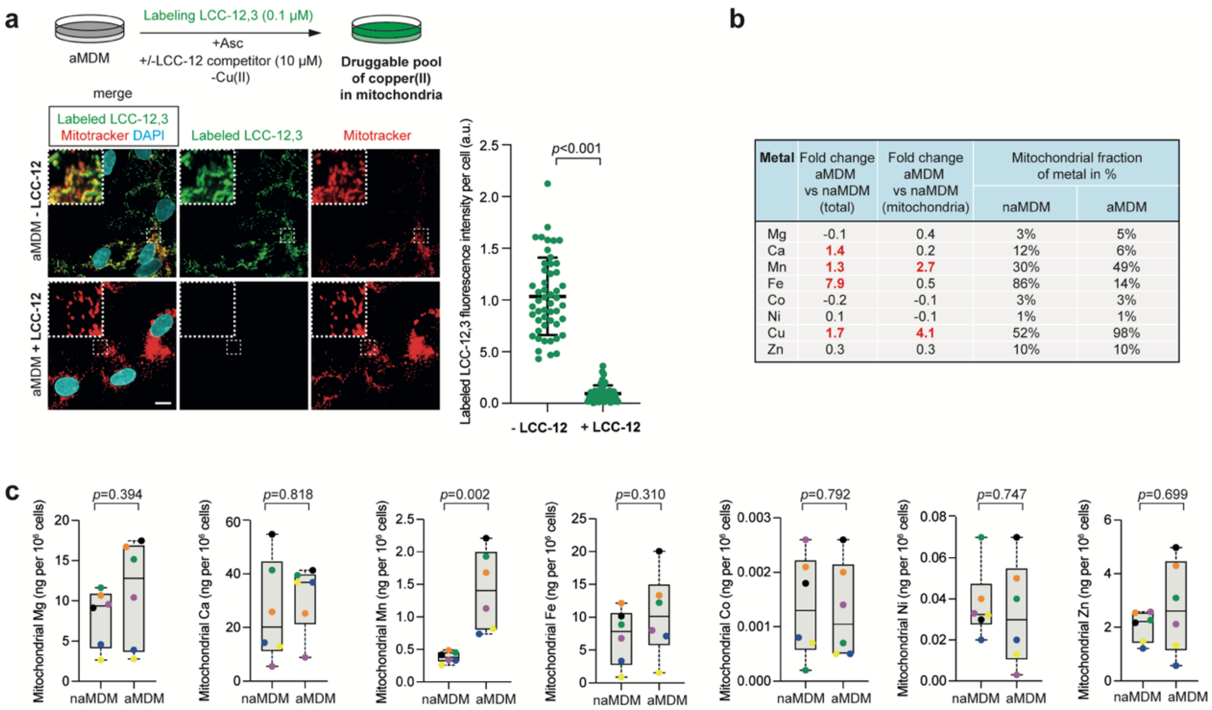


| Metal | Fold change aMDM vs naMDM (total) | Fold change aMDM vs naMDM (mitochondria) | Mitochondrial fraction of metal in % | |
|---|---|---|---|---|
| | | | naMDM | aMDM |
| Mg | -0.1 | 0.4 | 3% | 5% |
| Ca | 1.4 | 0.2 | 12% | 6% |
| Mn | 1.3 | 2.7 | 30% | 49% |
| Fe | 7.9 | 0.5 | 86% | 14% |
| Co | -0.2 | -0.1 | 3% | 3% |
| Ni | 0.1 | -0.1 | 1% | 1% |
| Cu | 1.7 | 4.1 | 52% | 98% |
| Zn | 0.3 | 0.3 | 10% | 10% |

**Extended Data Figure 3 | Detection of a druggable pool of copper(II) in mitochondria. a**, Fluorescence microcopy images of labeled LCC-12,3 (0.1 µM). Click labeling performed with or without LCC-12 competitor (10 µM) in absence of added copper in aMDM. Scale bar, 10 µm. Student's T-test. Mean ± SEM. At least 50 cells were quantified per condition. **b**, Comparison of total metal contents in cells and mitochondria of naMDM versus aMDM determined by ICP-MS. **c**, ICP-MS of mitochondrial metals in naMDM and aMDM. $n$=6 donors. Mann-Whitney test. Box plots with median and whiskers of lowest and highest values.

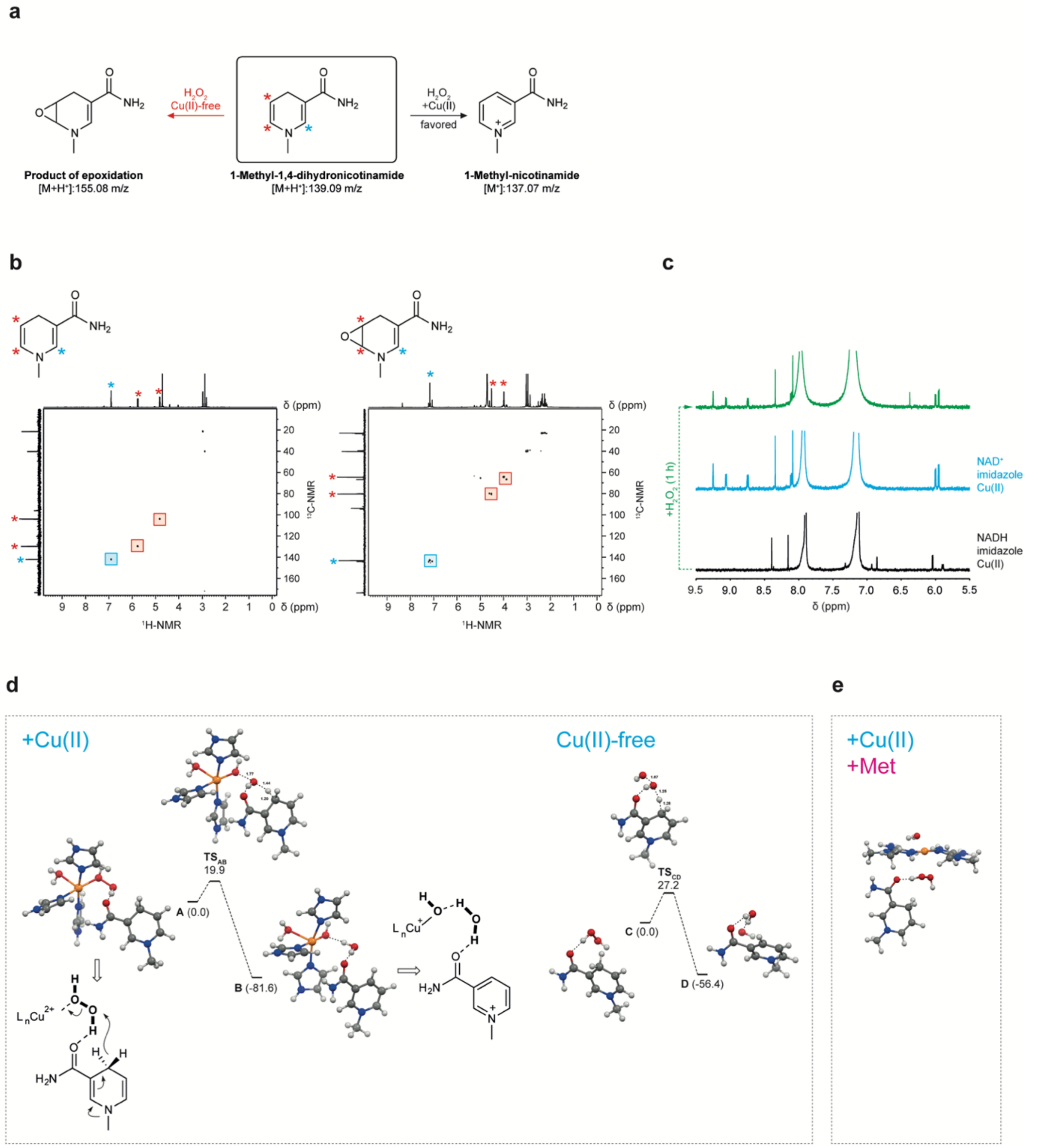


**Extended Data Figure 4 | Mitochondrial copper(II) regulates NAD(H) redox cycling. a**, Reaction of 1-methyl-1,4-dihydronicotinamide with hydrogen peroxide to afford either 1-methyl-nicotinamide or a product of epoxidation. The mass of molecular ions detected by mass spectrometry are indicated. **b**, Heteronuclear single quantum coherence (HSQC) NMR spectra of 1-methyl-1,4-dihydronicotinamide and its product of epoxidation. Red stars indicate $^{1}$H and $^{13}$C signals of the most reactive double bond towards hydrogen peroxide and that of the corresponding epoxide product. Blue stars mark hydrogens and carbons of the least reactive double bond towards hydrogen peroxide. Blue boxes show $^{1}$H-$^{13}$C HSQC correlations of the least reactive double bond. Red boxes show $^{1}$H-$^{13}$C HSQC correlations of the most reactive double bond and corresponding epoxide. **c**, $^{1}$H-NMR spectra of NADH in the presence of imidazole and copper(II) (black), NAD$^{+}$ in the presence of imidazole and copper(II) (blue), and the reaction product of NADH with hydrogen peroxide obtained in the presence of imidazole and copper(II) at 37 °C for 1 h (red). **d**, Free energy profile ($\Delta G_{298}$, kcal/mol) of the

[(Imidazole)$_3$Cu($H_2O$)](II)-mediated hydride-transfer reaction from 1-methyl-1,4-dihydronicotinamide to $H_2O_2$. Selected distances in Å. Free energy profile ($\Delta G_{298}$, kcal/mol) of the copper-free H-transfer reaction from 1-methyl-1-methyl-1,4-dihydronicotinamide to $H_2O_2$. Selected distances in Å. **e**, Optimized adduct of *syn* (metformin)$_2$Cu(II), 1-methyl-1,4-dihydronicotinamide, $H_2O_2$ and $H_2O$.

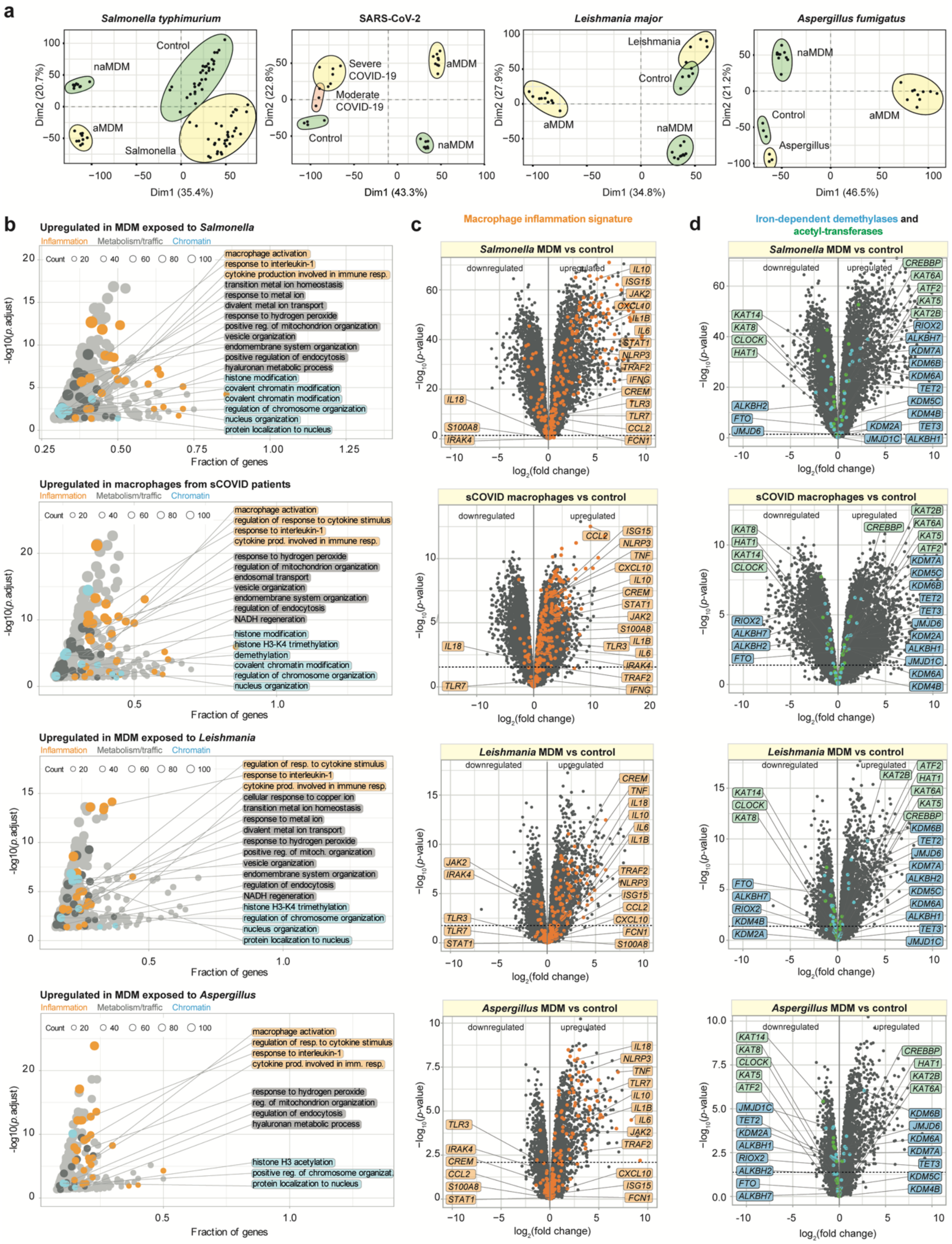


**Extended Data Figure 5 | Mitochondrial copper(II) regulates the epigenetic state and transcriptional programs. a**, Principal Component Analysis (PCA) of RNA-seq comparing naMDM (*n*=10 donors) and aMDM (*n*=10 donors) with MDM exposed to *Salmonella typhimurium*, macrophages from bronchoalveolar fluids of moderate and severe COVID-19 patients, MDM exposed to *Leishmania major* and MDM exposed to *Aspergillus fumigatus*. **b**, GO term analyses of upregulated genes in MDM exposed to *Salmonella typhimurium* versus control MDM, sCovid versus control macrophages, MDM exposed to *Leishmania major* versus control MDM and MDM exposed to *Aspergillus fumigatus* versus control MDM. **c**, RNA-seq analysis of gene expression in MDM exposed to *Salmonella typhimurium* versus control MDM, sCovid versus control macrophages, MDM exposed to *Leishmania major* versus control MDM and MDM exposed to *Aspergillus fumigatus* versus control MDM. Inflammatory gene signature is illustrated (orange). Dashed lines, adjusted *p*-value=0.05. **d**, RNA-seq analysis of gene expression in MDM exposed to *Salmonella typhimurium* versus control MDM, sCovid versus control macrophages, MDM exposed to *Leishmania major* versus control MDM and MDM exposed to *Aspergillus fumigatus* versus control MDM. Genes encoding iron-dependent demethylases (blue) and acetyl-transferases (green) are illustrated. Dashed lines, adjusted *p*-value=0.05.

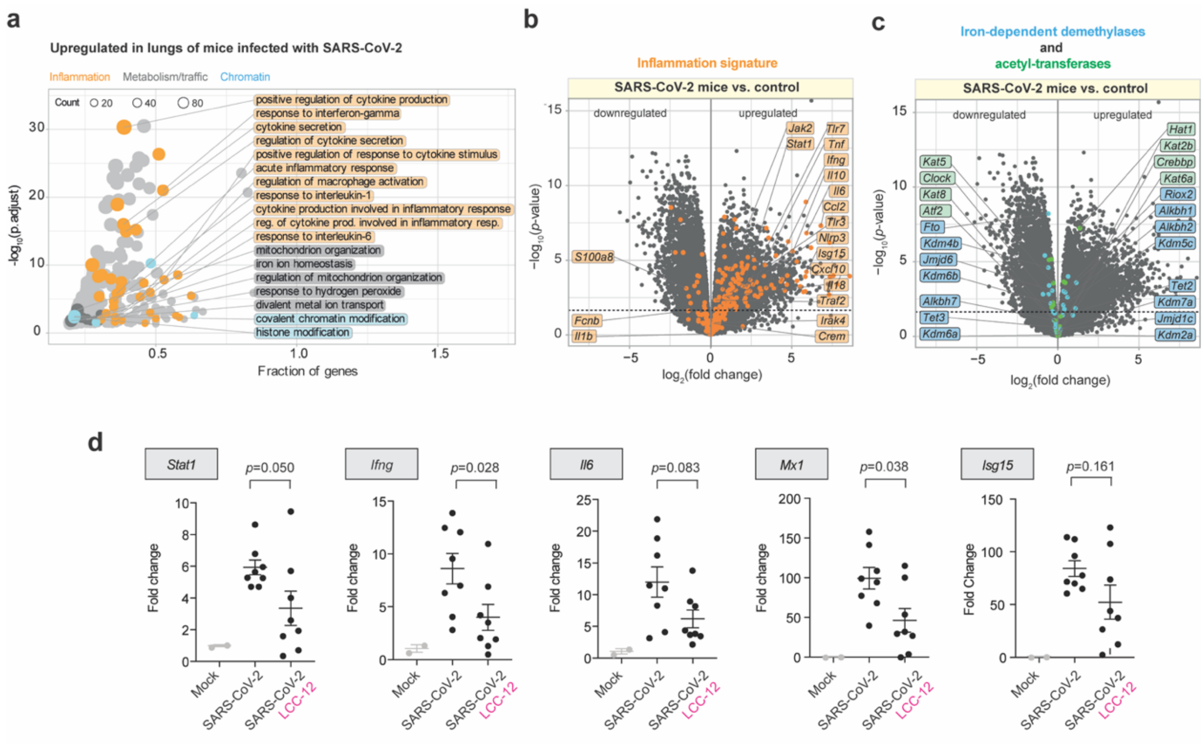


**Extended Data Figure 6 | Pharmacological inactivation of copper attenuates inflammation in a mouse models of viral infection.** **a**, GO term analysis of upregulated genes in the lungs of mice infected with SARS-CoV-2. **b**, RNA-seq analysis of gene expression in the lung tissues of mice infected with SARS-CoV-2. Inflammatory gene signature is illustrated (orange). Dashed lines, adjusted *p*-value=0.05. **c**, RNA-seq analysis of gene expression in the lungs of mice infected with SARS-CoV-2. Genes encoding iron-dependent demethylases (blue) and acetyl-transferases (green) are illustrated. Dashed lines, adjusted *p*-value=0.05. **d**, RT-qPCR analysis of inflammation markers in lung tissues of SARS-CoV2-infected mice treated with LCC-12 versus untreated. Mann-Whitney test. Mean values ± SEM.

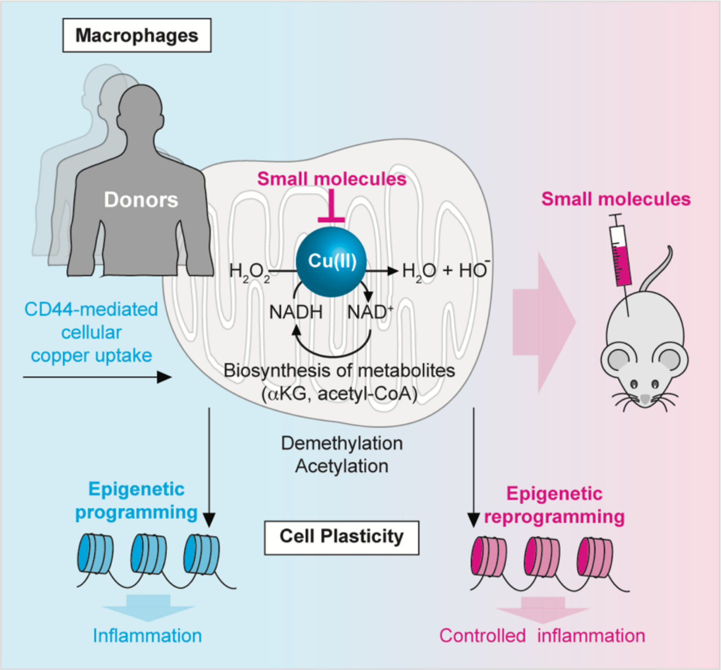


**Extended Data Figure 7 | Copper-signaling regulates cell plasticity and drives inflammation.** Cell plasticity involves upregulation of the surface marker CD44, which mediates uptake of metal ions. In the presence of copper(II), NADH reacts with hydrogen peroxide to replenish $NAD^+$, a key cofactor used for the biosynthesis of αKG and acetyl-CoA. These co-substrates of iron-dependent demethylases and acetyl transferases are required for epigenetic and transcriptional reprogramming. Pharmacological inactivation of mitochondrial copper(II) blocks NAD(H) redox cycling, leading to distinct epigenetic states and transcriptional profiles. Targeting copper(II) interfere with cell plasticity in immune and cancer cells, conferring therapeutic benefits.

**Captions for Supplementary Tables 1-9**

**Supplementary Table 1 |** Quantitative metabolomics analysis of mitochondrial cell extracts

**Supplementary Table 2|** Quantitative metabolomics analysis of total cell extracts

**Supplementary Table 3 |** RNA-seq analysis of naMDM, aMDM and aMDM treated with LCC-12

**Supplementary Table 4 |** RNA-seq analysis of bronchoalveolar macrophages from patients infected with SARS-CoV-2 and macrophages exposed to *Salmonella typhimurium*, *Leishmania major* or *Aspergillus fumigatus*

**Supplementary Table 5 |** GO term analysis of genes upregulated in aMDM, in macrophages from sCOVID patients and MDM exposed to *Salmonella typhimurium*, *Leishmania major* or *Aspergillus fumigatus*

**Supplementary Table 6 |** GO term analysis of genes downregulated in aMDM treated LCC-12 versus aMDM

**Supplementary Table 7 |** GO term analysis of genes upregulated in lung tissues from SARS-CoV-2-infected mice

**Supplementary Table 8 |** RNA-seq analysis of lung tissues from SARS-Cov-2-infected mice treated with LCC-12

**Supplementary Table 9 |** GO term analysis of genes downregulated in lung tissues from SARS-CoV-2-infected mice treated with LCC-12

### Materials and Methods

***Materials availability:*** All in-house reagents are available from the Lead Contact (R.R.) under a material transfer agreement with Institut Curie.

***Primary cells:*** Peripheral blood samples were collected from 80 healthy donors (Etablissement Français du Sang). Pan monocytes were isolated by negative magnetic sorting using microbeads according to the manufacturer's instructions (Miltenyi Biotec, 130-096-537) and cultured immediately without freezing in the presence of cytokines, to trigger *in vitro* differentiation as described in the method details.

***In vivo animal studies:*** Survival assessment using the LPS mouse model was conducted at Fidelta Ltd according to 2010/63/EU and National legislation regulating the use of laboratory animals in scientific research and for other purposes (Official Gazette 55/13). An institutional committee on animal research ethics (CARE-Zg) oversaw that animal-related procedures were not compromising the animal welfare. Experiments were performed on 8-week-old male BALB/c mice. Male littermates were randomly assigned to experimental groups. Flow cytometry and ICP-MS using the LPS mouse models were performed in accordance with French laws concerning animal experimentation (#202107221634651l) and approved by Institutional Animal Care and Use Committee (C2EA-47). Experiments were performed on 8-week-old male BALB/c mice and 5-week-old male SWISS mice. Male littermates were randomly assigned to experimental groups. All animal work using the CLP model was conducted in accordance with 2010/63/EU and National legislation regulating the use of laboratory animals in scientific research and for other purposes (Official Gazette 55/13). Experiments were performed on 9-week-old male BALB/c mice. An Institutional Committee on Animal Research Ethics (CEEA - 047) oversaw that animal-related procedures were not compromising the animal welfare. All animal work using the Sars-Cov2 model was conducted in accordance with 2010/63/EU and National legislation regulating the use of laboratory animals in scientific research and for other purposes (Official Gazette 55/13). An Institutional Committee on Animal Research Ethics (CEEA - 047) oversees that animal-related procedures are not compromising the animal welfare. Experiments were performed on 9-week-old male BALB/c mice.

***Antibodies:*** WB: Western blot, FC: flow cytometry, FM: fluorescence microscopy, IS: immunohistochemical staining, NS: NanoSIMS, CS: Cell sorting
*Anti-human antibodies:* CD14-Krome Orange (Beckman Coulter, B01175, FC), CD16-Pacific Blue (Beckman Coulter, A82792, FC), CD44 (Abcam, ab189524, WB), CD44-Alexa-Fluor-647 (Novus Biologicals, NB500-481AF647, FC), CD80-Alexa-Fluor-700 (Becton, Dickinson and Company, 561133, FC), CD86-PE/Cy7 (Becton, Dickinson and Company, 561128, FC), CD163-PE (Becton, Dickinson and Company, 556018, FC), Copper transporter 1 (Ctr1, Novus Biologicals, NB100-402AF647, FC), Copper transporter 1 (Ctr1, Abcam, ab129067, WB), Cytochrome *c* (Cyt *c*, Cell Signaling, 12963S, FM, NS), H3 (Cell Signaling, 9715S, FM), H3K4me3 (Diagenode, C15410003-50, FM), H3K9me2 (Cell Signaling, 4658S, FM), H3K9me3 (Cell Signaling, 13969S, FM), H3K27me3 (Cell Signaling, 9733S, FM), H3K36me2 (Abcam, ab9049, FM), H3K9ac (Cell Signaling, 9649S, FM), H3K14ac (Cell Signaling, 7627S, FM), H3K27ac (Cell Signaling, 8173S, FM), Superoxide dismutase 2 (SOD2, Abcam, ab13534, FM, WB), TfR1-APC-Alexa-Fluor-750 (Beckman Coulter, A89313, FC), Transferrin receptor 1 (TfR1, Life Technologies, 13-6800, WB), γ-Tubulin (Sigma-Aldrich, T5326, WB), Alexa-Fluor-488 anti-rabbit (Life Technologies, A-11008, FM), Alexa-Fluor-594 anti-mouse (Life Technologies, A-11032, FM), 10 nM gold-nanoparticle-loaded anti-rabbit (Abcam, ab27234, NS).
*Anti-mouse Antibodies:* CD11b-Pacific Blue (BioLegend, 101224, FC, CS), CD40-APC (BioLegend, 124612, FC), CD44 (Abcam, 189524, IS), CD44-AF488 (BioLegend, 103015, FC), CD45-BV510 (BioLegend, 103138, FC), CD71-BV711 (BD Biosciences, 740667, FC), CD86-PE (Biolegend, 105007, FC), CD170-PEeFluor 610 (eBioscience, 61-1702-80, FC), CD206-BV650 (BioLegend, 141723, FC), F4/80-PE (TONBO, TNB50-4801-U100, CS), F4/80-BV605 (BioLegend, 123133, FC), I-A/I-E-AF700 (BioLegend, 107622, FC), Ly6C-PerCP/Cy5.5 (BioLegend, 128012, FC, CS), Ly6G-AF647 (BioLegend, 127610, CS), Ly6G-PE/CY7 (BioLegend, 127618, FC), NOS2-APC (eBioscience, 17-5920-82, FC).

***Cell culture****:* This study was carried out using primary monocytes obtained from peripheral blood samples of 80 independent human donors (Etablissement Français du Sang). Pan monocytes were isolated by negative magnetic sorting using microbeads according to the manufacturer's instructions (Miltenyi Biotec, 130-096-537), and cultured in RPMI 1640 supplemented with GlutaMAX (Thermo Fisher Scientific, 11554516), 10% fetal bovine serum (FBS, CVFSVF00-01) and treated with granulocyte-macrophage colony-stimulating factor (GM-CSF, Miltenyi Biotec, 130-093-866, 100 ng/mL) to induce differentiation into monocyte-derived macrophages (MDM). At day 5 of differentiation, non-activated MDM (naMDM) were treated with lipopolysaccharides (LPS, InvivoGen, tlrl-3pelps, 100 ng/mL, 24 h) and interferon-γ (IFNγ, Miltenyi Biotec, 130-096-484, 20 ng/mL, 24 h) to generate activated MDM (aMDM). aMDM were cotreated with metformin (Met, 1,1-dimethylbiguanide hydrochloride, Alfa Aesar, J63361, 10 mM, 24 h), LCC-12 (in-house, 10 µM, 24 h), LCC-12,3 (in-house, 0.1 µM, 3 h), $^{15}$N-$^{13}$C-LCC-12 (in-house, 10 µM, 3 h, for NS), LCC-4,4 (in-house, 10 µM, 24 h), CCCP (Carbonyl cyanide 3-chlorophenylhydrazone, Sigma-Aldrich, C2759, 10 µM, 3 h), D-penicillamine (D-Pen, Sigma Aldrich, P4875, 250 µM, 24 h), Ammonium tetrathiomolybdate (ATTM, Sigma-Aldrich, 323446, 1 µM, 24 h) or anti-human CD44 therapeutic antibody (RG7356, Creative Biolabs, TAB-128CL, 10 µg/mL, 24 h) as indicated. As a comparison, to generate anti-inflammatory macrophages, pan monocytes were treated with macrophage colony-stimulating factor (M-CSF, Miltenyi Biotec, 130-096-492, 100 ng/mL), and IL-4 (Miltenyi Biotec, 130-093-921, 20 ng/mL) was added at day 5 for an additional day. CD4$^+$ lymphocytes were isolated from peripheral blood samples by negative magnetic sorting using microbeads according to the manufacturer's instructions (Miltenyi Biotec, 130-096-533) and cultured in RPMI1640 supplemented with glutamine and 10% fetal bovine serum. CD4$^+$ lymphocytes were activated for 48 h using CD3/CD28 antibodies (2,5 µg/mL), in presence of LCC-12 (10 µM). The activation status of lymphocytes was assessed by measuring CD25 and CD69 cell surface markers by flow cytometry. CD8$^+$ lymphocytes were isolated from peripheral blood samples by negative magnetic sorting using microbeads according to the manufacturer's instructions (Miltenyi Biotec, 130-096-495) and cultured in RPMI1640 supplemented with glutamine and 10% fetal bovine serum. CD8$^+$ lymphocytes were activated for 48 h using CD3/CD28 antibodies (2,5 µg/mL) and treated with LCC-12 (10 µM). The activation status of lymphocytes was assessed by measuring CD25 and CD69 cell surface markers by flow cytometry. For neutrophils, peripheral blood samples were collected and red cells in whole blood samples were lysed (ebioscience RBC lysis buffer, 00-4300-54). The remaining cells were cultured in RPMI 1640 supplemented with glutamine, and 2% human serum, and activated 1 h with LPS (2 µg/mL) in presence of LCC-12 (10 µM). The neutrophil population was determined by flow cytometry using FSC, SSC and CD15 surface markers. The activation status of neutrophils was assessed by measuring CD64 and CD66b cell surface markers by flow cytometry. Monocytes were isolated from peripheral blood samples by negative magnetic sorting using microbeads according to the manufacturer's instructions (Miltenyi Biotec, 130-096-537), and cultured in RPMI 1640 supplemented with glutamine, 10% fetal bovine serum. Monocytes were treated with lipopolysaccharides (LPS, 100 ng/µL, 24 h) to generate activated monocytes and these were cotreated with LCC-12 (in-house, 10 µM, 24 h). The activation status of monocytes was assessed by measuring CD25 and CD80 cell surface markers by flow cytometry. For dendritic cells, pan monocytes were isolated by negative magnetic sorting using microbeads according to the manufacturer's instructions (Miltenyi Biotec, 130-096-537), and cultured in RPMI 1640 supplemented with glutamine, 10% fetal bovine serum and treated with granulocyte-macrophage colony-stimulating factor (GM-CSF, 100 ng/mL) and IL-4 (10 ng/mL) to induce differentiation into dendritic cells (DC). At day 5 of differentiation, DC were treated with lipopolysaccharides (LPS, 100 ng/µL, 24 h) to generate activated DC and were cotreated with LCC-12 (in-house, 10 µM, 24 h). The activation status of the dendritic cells was assessed by measuring CD40, CD83, CD80 and CD86 cell surface markers by flow cytometry. Mouse pancreatic cancer cells FC1245 were cultured in Dulbecco's Modified Eagle Medium GlutaMAX (DMEM, ThermoFisher Scientific, 61965059) supplemented with 10% FBS (Gibco, 10270-106) and Penicillin-Streptomycin mixture (BioWhittaker/Lonza, DE17-602E). Cells were treated with transforming growth factor ß (TGF-ß, Miltenyi Biotec, 130-095-066, 10 ng/mL) for 3 d.

***Flow cytometry:*** Cells were washed with ice-cold 1× PBS, incubated with Fc block (Human TruStain FcX, Biolegend, 422302, 1:20) for 15 min, incubated with antibodies for 20 min at 4 °C and washed before analysis using a flow cytometer (BD LSRFortessa X-20). Macrophages were analyzed with an antibody panel consisting of antibodies against the following cell surface proteins: CD14, CD16, CD44, CD80, CD86, CD163, Ctr1 and TfR1. The data were analyzed with FlowJo software v. 10.0.00003. *Flow cytometry analysis of mitochondrial $H_2O_2$ content:* cells were incubated with MitoPY1 (R&D Systems, 4428/10, 5 μM, 24 h) during activation with LPS and IFNγ after day 5 of differentiation with GM-CSF. Fluorescence was analyzed by flow cytometry.

***Inductively coupled plasma mass spectrometry (ICP-MS):*** HA (Carbosynth, FH45321, 600-1000 kDa, 1 mg/mL) or the anti-CD44 antibody RG7356 was added together with LPS and IFNγ and cells were treated for 24 h. Glass vials equipped with Teflon septa were cleaned with nitric acid 65% (VWR, Suprapur, 1.00441.0250), washed with ultrapure water (Sigma-Aldrich, 1012620500) and dried. MDM and aMDM were harvested followed by two washes with 1× PBS. Cells were then counted using an automated cell counter (Entek) and transferred in 200 μL 1× PBS to the cleaned glass vials. Mitochondria were extracted as described in the mitochondrial extraction section from a pre-counted population of cells. Samples were lyophilized using a freeze dryer (CHRIST, 22080). 200 μL 1× PBS were used as a control in duplicate per condition. For tumor and healthy tissue samples, small pieces were cut and dried in glass vials and weighed. Final results for these samples are normalized against dry weight. Samples were subsequently mixed with nitric acid 65% overnight and heated at 80 °C for 2 h. Samples were diluted with ultrapure water to a final concentration of 0.475 N nitric acid and transferred to metal-free centrifuge vials (VWR, 89049-172) for subsequent ICP-MS analysis. Amounts of metals were measured using an Agilent 7900 ICP-QMS in low-resolution mode, taking natural isotope distribution into account. Sample introduction was achieved with a micro-nebulizer (MicroMist, 0.2 mL/min) through a Scott spray chamber. Isotopes were measured using a collision-reaction interface with helium gas (5 mL/min) to remove polyatomic interferences. Scandium and indium internal standards were injected after inline mixing with the samples to control the absence of signal drift and matrix effects. A mix of certified standards was measured at concentrations spanning those of the samples to convert count measurements to concentrations in the solution. Uncertainties on sample concentrations were calculated using algebraic propagation of ICP-MS blank and sample counts uncertainties. Values were normalized against cell number or dry weight.

***siRNA transfection:*** Human primary monocytes were transfected with Human Monocyte Nucleofector kit (Lonza,VPA-1007) according to the manufacturer's instructions. $5 \times 10^6$ monocytes were resuspended into 100 μL of nucleofector solution with 200 pmol of ON-TARGETplus CD44 SMARTpool siRNA (Horizon Discovery, L-009999-00-0050) or negative control siRNA (Qiagen, 1027310) before nucleofection with Nucleofector II (Lonza). Cells were then immediately removed and incubated overnight with 5 mL of prewarmed complete RPMI medium (Thermo Fisher Scientific). The following day, GM-CSF was added to the medium.

***Western blotting:*** Cells were treated as indicated and then washed with 1× PBS. Proteins were solubilized in 2× Laemmli buffer containing benzonase (VWR, 70664-3, 1:100). Extracts were incubated at 37 °C for 1 h, and quantified using a NanoDrop 2000 spectrophotometer (Thermo Fisher Scientific). Protein lysates were resolved by SDS-PAGE electrophoresis (Invitrogen sure-lock system and Nu-PAGE 4–12% Bis-Tris precast gels) and transferred onto nitrocellulose membranes (Amersham Protran 0.45 μm) using a Trans-Blot SD semi-dry electrophoretic transfer cell (Bio-rad). Membranes were blocked with 5% non-fat skimmed milk powder in 0.1% Tween-20/1× PBS for 1 h. Blots were then probed with the relevant primary antibodies in 5% BSA, 0.1% Tween-20/1× PBS at 4 °C overnight with gentle motion. Membranes were washed with 0.1% Tween-20/1× PBS three times and incubated with horseradish peroxidase conjugated secondary antibodies (Jackson Laboratories) in 5% non-fat skimmed milk powder, 0.1% Tween-20/1× PBS for 1 h at room temperature and washed three times with 0.1% Tween-20/1× PBS. Antigens were detected using the SuperSignal West Pico PLUS chemiluminescent detection kits (Thermo Fisher Scientific, 34580 and 34096). Signals were recorded using a Fusion Solo S Imaging System (Vilber) and quantified with ImageJ using pixel intensity normalized against the signal of γ-tubulin.

***Fluorescence microscopy:*** Isolated monocytes were plated on coverslips, differentiated and activated as described in the cell culture paragraph. For fluorescent detection of HA and lysosomal Fe(II) or Cu(II), live cells were treated with HA-FITC (800 kDa, Carbosynth, YH45321, 0.1 mg/mL, ~125 μM) and Lys-Cu (in-house, 20 μM, 1 h)[51] for 1 h before fixation. Cells were washed three times with 1× PBS, fixed with 2% paraformaldehyde in 1× PBS for 12 min and then washed three times with 1× PBS. For antibody staining, cells were then permeabilized with 0.1% Triton X-100 in 1× PBS for 5 min and washed three times with 1× PBS. Subsequently, cells were blocked in 2% BSA, 0.2% Tween-20/1× PBS (blocking buffer) for 20 min at room temperature. Cells were incubated with the relevant antibody in blocking buffer for 1 h at room temperature, washed three times with 1× PBS and were incubated with secondary antibodies for 1 h. Finally, coverslips were washed three times with 1× PBS and mounted using VECTASHIELD containing DAPI (Vector Laboratories, H-1200-10). Fluorescence images were acquired using a Deltavision real-time microscope (Applied Precision). 40×/1.4NA, 60×/1.4NA and 100×/1.4NA objectives were used for acquisitions and all images were acquired as z-stacks. Images were deconvoluted with SoftWorx (Ratio conservative - 15 iterations, Applied Precision) and processed with ImageJ. Colocalization quantification was calculated using ImageJ. Histone quantification was performed using ImageJ by delineating the nuclei using DAPI fluorescence, and calculating the mean fluorescence intensity normalized by area.

***UV titration experiments:*** To a solution of LCC-12 (5 μM) in HEPES (10mM), portions of 0.1 mol equiv. of a solution of $CuCl_2$ in HEPES (10 mM) were added up to 3 mol equiv. UV spectra were recorded on an Analytik Jena UV/ VIS spectrophotometer specord 205 system at room temperature in the 200-1000 nm range using a micro cuvette (quartz Excellence Q 10 mm). All spectra were blanked against HEPES buffer.

***Bright-field microscopy and digital photographs:*** Bright field images were acquired using a CKX41 microscope (Olympus) and cellSens Entry imaging software (Olympus). Digital images were taken with an iPhone 11 Pro (Apple).

***NMR spectroscopy:*** $^{1}H$-NMR, $^{13}C$-NMR and $^{15}N$-NMR spectra were recorded on a 500 MHz Bruker spectrometer at 310 K, and chemical shifts δ are expressed in ppm using the residual non-deuterated solvent signal as internal standard. To measure the interaction of HA tetrasaccharide with copper, portions of 0.25 mol equiv. of a solution of $CuCl_2$ in $D_2O$ (8.6 mg in 599 μL $D_2O$) were added to a 2 mM solution of HA tetrasaccharide (TCI Chemicals, H1284) in $D_2O$ (1.0 mg HA tetrasaccharide in 600 μL $D_2O$) up to 1 mol equiv. into an NMR tube. Then, a drop of trifluoroacetic acid (TFA, Alfa Aesar, A12198) was added. In a separate NMR tube, a drop of TFA was added to a copper-free 2 mM solution of HA tetrasaccharide in $D_2O$. To control measure oxidation of NADH into $NAD^+$, $^{1}H$-NMR were recorded and tubes were prepared as follows: β-Nicotinamide adenine dinucleotide, reduced (NADH, Sigma-Aldrich, N4505-100MG, 200 μM) or β-Nicotinamide adenine dinucleotide ($NAD^+$, Sigma Aldrich, N0632-1G, 200 μM), imidazole (10 mM), $CuSO_4$ (10 μM) were added to an NMR tube containing sodium phosphate buffer in $D_2O$ (10 mM, pD = 8.4, 981 μL) and $^1H$-NMR were recorded at $t_0$. $H_2O_2$ (19 μL of 100 × diluted solution of 32.3 % wt in $H_2O$) was added to the NMR tubes and $^1H$-NMR were recorded after 1 h. To measure the interaction of $^{15}N$-$^{13}C$-LCC-12 with copper, 1 mol equiv. of a solution of $CuCl_2$ in $D_2O$ (10.0 mg in 70.3 μL $D_2O$) were added to a 10 mM solution of $^{15}N$-$^{13}C$-LCC-12 in $D_2O$ (3.0 mg in 600 μL $D_2O$) into an NMR tube. Then 2 mol equiv. of $H_2SO_4$ (Prolabo VWR, 20700.298) were added. In a separate NMR tube containing 10 mM of $^{15}N$-$^{13}C$-LCC-12 in $D_2O$, 2 mol equiv. of $H_2SO_4$ were added NMR spectra were recorded immediately after addition of each component.

***Chemical synthesis:*** Products were purified on a preparative HPLC Quaternary Gradient 2545 equipped with a Photodiode Array detector (Waters) fitted with a reverse phase column (XBridge Prep C18 5μm OBD 30×150 mm). NMR Spectra were run in DMSO-$d_6$ or Methanol-$d_6$ at 298 K unless stated otherwise. $^1H$-NMR spectra were recorded on Bruker spectrometers at 400 or 500 MHz. Chemical shifts δ are expressed in ppm using the residual non-deuterated solvent signal as internal standard. The following abbreviations are used: ex, exchangeable; s, singlet; d, doublet; t, triplet; td, triplet of doublets; brs, broad signal; m, multiplet. The $^{13}C$-NMR spectra were recorded at 100.6 or 125.8 MHz, and chemical shifts δ are expressed in ppm using deuterated solvent signal as internal standard. The purity of final compounds, determined to be >98% by UPLC-MS, and low-resolution mass spectra (LRMS) were recorded on a Waters Acquity H-class equipped with a Photodiode array detector and SQ Detector 2 (UPLC-MS) fitted with a reverse phase column (Aquity UPLC BEH C18 1.7 μm, 2.1x50 mm). High resolution mass spectra (HRMS) were recorded on a Thermo Fisher Scientific Q-Exactive Plus equipped with a Robotic TriVersa NanoMate Advion.

*Lipophilic copper clamp 12 (LCC-12)*: Dicyandiamide (Alfa Aesar, A10451, 500 mg, 5.94 mmol), 1,12-diaminododecane (Alfa Aesar, A04258, 500 mg, 2.50 mmol) and $CuCl_2$ (Sigma-Aldrich, 22.201-1, 249 mg, 1.85 mmol) were suspended in water (6 mL) in a sealed tube and stirred for 1 h, then heated at 80 °C for 48 h. The resulting pink mixture was filtered and the solid was re-suspended in water (10 mL). $H_2S$, generated from dropwise addition of 37% aqueous (aq.) HCl (Supelco, 1.00317.100) on FeS (≃100 mesh powder, Alfa Aesar, 17422), was passed into the mixture until it turned black. The black mixture was filtered, and the filtrate was acidified to pH = 5 with a 1 M aq. solution of HCl. The solvent was evaporated under reduced pressure. LCC was purified by preparative HPLC ($H_2O$/$CH_3CN$/formic acid, 95:5:0.1 to 0:100:0.1) to give the LCC di-formic acid salt as a white powder (280 mg, 24 %). $^1H$-NMR (500 MHz, DMSO-$d_6$) δ: 8.80-8.08 (brs, 2H, ex), 8.47 (s, 2H, formate), 7.60-6.78 (brs, 12H, ex), 3.05 (brs, 4H), 1.43 (brs, 4H), 1.32-1.18 (m, 16H) ppm. $^{13}C$-NMR (125.8 MHz, DMSO-$d_6$) δ: 167.2 (formate), 160.4, 159.3, 41.3, 29.5 (3C), 29.3, 26.8 ppm. HRMS (ESI+) *m/z*: calculated for $C_{16}H_{38}N_{10}$ $[M+2H]^{2+}$ 185.1635, found 185.1637.

The synthesis of *bis-(cyanoguanidino)butane* was adapted from a previously published procedure[52]. 1,4-diaminobutane (112120250, Acros organics, 554 mg, 5,68 mmol) was dissolved in water and stirred with 37% aq. HCl for 10 min at room temperature (rt). The solvent was evaporated under reduced pressure and the resulting salt was suspended in butanol (5 mL) with sodium dicyanamide (178322, Sigma Aldrich, 1.09 g, 11.4 mmol) and stirred at 140 °C overnight. After filtration the solid was washed with butanol and cold water and recrystallized from water to give the bis-(cyanoguanidino)butane as a white powder (350 mg, 27 %). $^1H$-

NMR (500 MHz, DMSO-$d_6$) δ = 7.62-6.12 (m, 6H, ex), 3.03 (brs, 4H), 1.39 (brs, 4H) ppm. $^{13}$C-NMR (125.8 MHz, DMSO-$d_6$) δ = 161.6, 118.8, 40.9, 26.8 ppm.

*Lipophilic copper clamp 4,4 (LCC-4,4)*: Bis-(cyanoguanidino)butane (200 mg, 0.90 mmol) and buthylamine hydrochloride (B0710, TCI Chemical, 197 mg, 1.80 mmol) were mixed together in a sealed tube and heated at 150 °C without solvent for 4 h. After cooling to rt, the mixture was taken up in EtOH and a large excess of ethyl acetate (EtOAc) was added. The white precipitate was filtered and purified by preparative HPLC ($H_2O$/Acetonitrile/formic acid, 100:0:0.1 to 50:50:0.1) to give the LCC-4,4 di-formic acid salt as a white powder (130 mg, 31 %). $^1$H-NMR (400 MHz, DMSO-$d_6$) $\delta$ = 8.82-7.77 (m, 4H, ex), 8.46 (s, 2H, formate), 7.56-6.65 (brs, 8H, ex), 3.16-2.99 (m, 8H), 1.54-1.37 (m, 8H), 1.37-1.20 (m, 4H), 0.86 (t, $J$ = 6.8 Hz, 6H) ppm. $^{13}$C-NMR (100.6 MHz, DMSO-$d_6$) δ: 167.3 (formate), 159.1 (2C), 40.9 (2C), 31.6, 26.8, 20.0, 14.2 ppm. HRMS (ESI+) m/z: calculated for $C_{16}H_{38}N_{10}$ $[M+2H]^{2+}$ 185.1635, found 185.1636.

The synthesis of *bis-(cyanoguanidino)dodecane* was adapted from a previously published procedure [52]. 1,12-diaminododecane (500 mg, 2.5 mmol) was dissolved in a mixture of water and methanol and stirred with 37% aq. HCl for 10 min at rt. The solvent was evaporated under reduced pressure and the resulting salt was suspended in butanol (2.5 mL) with sodium dicyanamide (444 mg, 5.0 mmol) and stirred at 140 °C overnight. After filtration the solid was washed with butanol and cold water and recrystallized from a mixture of water/ethoxyethanol (2:1) to give the bis-(cyanoguanidino)dodecane as a white powder (365 mg, 44 %). $^1$H-NMR (400 MHz, DMSO-$d_6$) $\delta$ = 7.21-6.18 (m, 6H, ex), 3.03 (m, 4H), 1.40 (m, 4H), 1.33-1.14 (m, 16H) ppm. $^{13}$C-NMR (100.6 MHz, DMSO-$d_6$) $\delta$ = 161.6, 118.8, 41.0, 29.4 (3C), 29.2, 26.7 ppm.

*Lipophilic copper clamp 12,3 (LCC-12,3)*: Bis-(cyanoguanidino)dodecane (227 mg, 0.60 mmol) and but-3-yne-1-amine hydrochloride (EN300-76524, Enamine, 126 mg, 1.20 mmol) were mixed together in a sealed tube and heated at 150 °C without solvent for 4 h. After cooling to rt, the mixture was taken up in EtOH and a large excess of EtOAc was added slowly. The white precipitate was filtered and purified by preparative HPLC ($H_2O$/Acetonitrile/formic acid, 95:5:0.1 to 40:60:0.1) to give LCC-12,3 di-formic acid salt as a white powder (102 mg, 30 %). $^1$H-NMR (500 MHz, DMSO-$d_6$) $\delta$ = 9.05-7.60 (brs, 4H, ex), 8.47 (s, 2H, formate), 7.60-6.80 (m, 8H, ex), 3.29-3.16 (m, 4H), 3.06 (brs, 4H), 2.84 (s, 2H), 2.39-2.29 (m, 4H), 1.44 (brs, 4H), 1.25 (brs, 16H) ppm. $^{13}$C-NMR (125.8 MHz, DMSO-$d_6$) $\delta$ = 167.6 (formate), 159.7, 158.4, 82.6, 72.7, 41.3, 40.3, 29.5 (3C), 29.2, 26.8, 19.4 ppm. HRMS (ESI+) *m/z*: calculated for $C_{24}H_{46}N_{10}$ $[M+2H]^{2+}$ 237.1948, found 237.1947.

*Isotopically labelled lipophilic copper clamp*: Dicyandiamide $^{15}$N and $^{13}$C marked (Eurisotop, CNLM-9324-PK, 50 mg, 0.55 mmol), 1,12-diaminododecane (46.8 mg, 0.23 mmol) and $CuCl_2$ (31.4 mg, 0.23 mmol) were suspended in a sealed tube in water (0.6 mL) and stirred for 1 h, then heated at 80 °C for 48 h. The resulting pink mixture was acidified with an aq. solution of HCl (2 M, 1 mL) until complete dissolution of the precipitate. The mixture was concentrated under reduced pressure and isotopically labeled LCC-12 was purified by preparative HPLC ($H_2O$/Acetonitrile/formic acid, 95:5:0.1 to 73:27:0.1) to give the isotopically labeled LCC-12 di-formic acid salt as a white powder (35 mg, 32 %). $^1$H-NMR (500 MHz, DMSO-$d_6$) δ: 8.60-7.76 (brs, 2H, ex), 8.48 (s, 2H, formate), 7.70-6.50 (brs, 12H, ex), 3.05 (brs, 4H), 1.43 (m, 4H), 1.33-1.18 (m, 16H) ppm. $^{13}$C-NMR (125.8 MHz, DMSO-$d_6$) δ: 167.0 (formate), 160.2, 159.2, 41.3, 29.5 (3C), 29.2, 26.8 ppm.

***Structural characterization of metformin-based Cu complexes:*** The starting structure for Cu(metformin)$_2$ was based on the published X-ray structure[53]. The starting geometries for the other copper(II) complexes were obtained via molecular dynamics conformation search (Gabedit[54], amber99[55] potential). For each complex, the 10 geometries with the lowest energies resulting from the MD search were reoptimized using MOPAC2016 (PM7[56], COSMO[57] water model). The geometries with the lowest energy for each complex were optimized at TPSSh/D3BJ/Def2-TZVP level using Orca 4.2.1[58]. We performed a benchmark study based on the structure of Cu(metformin)$_2$ using B3LYP[59], M062X[60], TPSSh, BHLYP[61,62] functionals with the Def2-TZVP[63] basis set using D3BJ[64] dispersion correction and the CPCM water solvation model. We have also used the BHLYP functional with the SVP basis set and the SMD water solvation model, which was recommended in the literature[65] for copper(II) complexes.

***High-resolution mass spectrometry of biguanide:metal complexes:*** HRMS solutions were prepared and injected without further dilution. Stock solution of LCC-12·di-formate/$K_2CO_3$ (1:2) (100 µM) or metformin·HCl/$K_2CO_3$ (1:1) (200 µM) were prepared in analytical grade methanol. Stock solutions of metals, $CuCl_2 \cdot 2H_2O$ (405840050, Acros Organics, 100 µM), $MnCl_2$ (244589, Sigma-Aldrich, 100 µM), $CaCl_2 \cdot 2H_2O$ (423520250, Acros Organics, 100 µM), $FeCl_2 \cdot xH_2O$ (12357, Alfa Aesar, 100 µM), $MgCl_2 \cdot 6H_2O$ (25 108.295, Prolabo, 100 µM), $NiCl_2$ (53131, Alfa Aesar, 100 µM) and $ZnCl_2$ (429430, Sigma-Aldrich, 100 µM) were prepared in milliQ water. The HRMS solutions were prepared in methanol (80 µL) with LCC-12·di-formate/$K_2CO_3$ (10 µL) and the relevant metal (10 µL) with a 1:1 ratio, or with metformin·HCl/$K_2CO_3$ (10 µL) and $CuCl_2 \cdot 2H_2O$ (10 µL) with a 2:1 ratio.

***Nanoscale secondary ion mass spectrometry (NanoSIMS):*** MDM were grown on coverslips and activated to obtain aMDM as described in the cell culture section. Cells were treated with 10 µM $^{15}N$-$^{13}C$-LCC-12 for 3 h. Subsequently, cells were washed twice with 1× PBS, once with 0.1 M cacodylate buffer (LFG Distribution, 11653) and then fixed with 2% paraformaldehyde in 0.1 M cacodylate buffer for 20 min. Then, cells were washed three times with 0.1 M cacodylate buffer for 5 min and permeabilized with 0.1% Triton-X in 0.1 M cacodylate buffer for 5 min. Subsequently, cells were washed three times with 0.1 M cacodylate buffer and blocking buffer (2% BSA, 0.1% Tween in 0.1 M cacodylate buffer) was added for 20 min. Primary antibody (1:400) was added for 1 h in blocking buffer. Then, cells were washed three times with 0.1 M cacodylate buffer and the 10 nM gold-nanoparticle-loaded secondary antibody (1:50) was added in blocking buffer for 1 h. Cells were washed three times with 0.1 M cacodylate buffer and treated with 1% $OsO_4$ (Electron Microscopy Sciences, 19152) in 0.1 M cacodylate buffer for 1 h. Coverslips with samples were washed three times for 10 min with Milli-Q water. Subsequently, cells were dehydrated with sequential EtOH solutions each for 10 min each: 50%, 70%, 2 × 90%, 3× 100% (dried over molecular sieves, Sigma-Aldrich, 69833). Samples were then coated with a 1:1 mixture of resin (Electron Microscopy Sciences, dodecenylsuccinic anhydride, 13710, methyl nadic anhydride, 19000, DMP-30, 13600 and LADD research industries: LX112 resin, 21310) and dry EtOH for 1 h. Then, samples were embedded in pure resin for 1 h. Embedding capsules (Electron Microscopy Sciences, 69910-10) were filled with resin, inverted onto the cover slides and placed in an oven at 56 °C for 24 h. 0.2 µm sections were prepared using a Leica Ultracut UCT microtome. Sample sections were deposited onto a clean silicon chip (Institute for Electronic Fundamentals/CNRS and University Paris Sud) and dried upon exposure to air before being introduced into the NanoSIMS-50 ion microprobe (Cameca, Gennevilliers, France). A $Cs^+$ primary ion was employed to generate negative secondary ion from the sample surface. The probe steps over the image field and the signal of selected secondary ion species were recorded pixel-by-pixel to create 2D images. Image of $^{12}C^{14}N^-$ was recorded to provide the anatomic structure of the cells, while the one of

$^{31}P^-$ highlights the location of cell nucleus. The cellular distribution of $^{15}N$-label was imaged by measuring the excess in $^{12}C^{15}N^-$ to $^{12}C^{14}N^-$ ratio with respect to the natural abundance level (0.0037), and the one for antibody with gold staining targeting mitochondria was performed by detecting directly $^{197}Au^-$ ion. When detecting $^{12}C^{15}N^-$ ion, appropriate mass resolution power was required to discriminate abundant $^{13}C^{14}N^-$ isobaric ions (with an M/ΔM of 4272). For each image recording process, multiframe acquisition mode was applied and hundreds of image planes were recorded. The overall acquisition time corresponding to the $^{15}N$ image was 12 h and 6 h 30 min for the $^{197}Au$ image. During image processing with ImageJ, the successive image planes were properly aligned using TomoJ plugin[66], so as to correct the slight primary beam shift during long hours of acquisition. A summed image was then obtained with improved statistics. Further, for the $^{12}C^{15}N^-$ to $^{12}C^{14}N^-$ ratio map, an HSI (Hue-Saturation-Intensity) color image was generated using OpenMIMS for display with increased significance[67]. The hue corresponds to the absolute $^{15}N/^{14}N$ ratio value, and the intensity at a given hue is an index of the statistical reliability.

***Click labeling:*** aMDM on coverslips were treated with LCC-12,3 (in-house, 0.1 µM, 3 h) in absence or presence of CCCP (10 µM, 3 h) fixed and permeabilized as indicated in fluorescence microscopy. Mitotracker was added to live cells for 45 min to before fixation. The click reaction cocktail was prepared using the Click-iT EdU Imaging kit (Life Technologies, C10337) according to the manufacturer's protocol. We mixed 430 µL of 1× Click-iT reaction buffer with 20 µL of $CuSO_4$ solution, 1.2 µL Alexa-Fluor-azide, 50 µL reaction buffer additive (sodium ascorbate) to reach a final volume of ~500 µL. Reactions were performed with or without $CuSO_4$ or ascorbate. Coverslips were incubated with the click reaction cocktail in the dark at room temperature for 30 min, then washed three times with 1× PBS. Immunofluorescence was then performed as described in fluorescence microscopy.

***Isolation of mitochondria:*** Mitochondria were isolated using the Qproteome Mitochondria Isolation Kit (Qiagen, 37612) according to the manufacturer's protocol. Cells were washed and centrifuged at 500× g for 10 min and the supernatant was removed. Cells were then washed with a solution of 0.9 % NaCl (Sigma-Aldrich, S7653-250G) and resuspended in ice-cold Lysis Buffer and incubated at 4 °C for 10 min. The lysate was then centrifuged at 1000× g for 10 min at 4 °C and the supernatant carefully removed. Subsequently, the cell pellet was resuspended in disruption buffer. Complete cell disruption was obtained by using a dounce homogenizer (mitochondria for ICP-MS) or a blunt-ended needle and a syringe (mitochondria for metabolomics). The lysate was then centrifuged at 1000× g for 10 min at 4 °C and the supernatant transferred to a clean tube. The supernatant was then centrifuged at 6000× g for 10 min at 4 °C to obtain mitochondrial pellets.

***Copper-catalyzed oxidation of NADH:*** The oxidation kinetics of NADH (N4505, Sigma Aldrich) was followed by measurement of absorbance at 340 nm using a Cary 300 UV-Vis spectrometer. The measures were recorded at 37 °C controlled with a Pelletier Cary temperature controller (Agilent Technologies). Stock solutions of NADH (1 mM), Imidazole (56750, Sigma Aldrich, 100 mM), $CuSO_4$ (451657, Sigma Aldrich, 500 µM), LCC-12 (10 mM or 1 mM) or metformin·HCl (J63361, Alfa Aesar, 100 mM or 10 mM) or LCC-4,4 (10 mM) were prepared in a 10 mM sodium phosphate buffer adjusted to pH = 8.0. The concentration of $H_2O_2$ (16911, Sigma Aldrich, 32.3 % wt in $H_2O$) was determined by titration with $KMnO_4$ and diluted 100 times in the phosphate buffer. The 1 mL experimental solutions were prepared in disposable cuvettes, with sodium phosphate buffer, NADH (200 µL, 200 µM), imidazole (100 µL, 10 mM), w/o $CuSO_4$ (20 µL, 10 µM), w/o LCC (20 µL, 20 µM or 200 µM), w/o metformin (40 µL, 400 µM or 4 mM) and $H_2O_2$ solution (19 µL, 2 mM) added at $t_0$. The concentration of

NADH was calculated from the measured absorbance at 340 nm and its molar extinction coefficient.

***Measurement of NADH concentrations:*** NADH absolute concentrations were measured using a fluorometric assay (Abcam, ab176723) according to the manufacturer's protocol. At least 500.000 cells were harvested per condition. Floating cells were harvested and adherent cells were washed with 1× PBS. Adherent cells were incubated with 1× PBS with 10 mM EDTA and then scraped and pooled together with the harvested floating cells. Cells were subsequently washed with ice-cold 1× PBS and counted. Cells were centrifuged at 1500 rpm for 5 min and the supernatant discarded. The pellet was then re-suspended in 100 µL lysis buffer (kit component) and incubated at 37 °C for 15 min. $NAD^+$ and NADH extraction solutions as well as $NAD^+$/NADH control solutions (kit components) were added and incubated at 37 °C for 15 min at a volume of 15 µL sample to 15 µL of the respective buffers (kit components). The reactions were stopped using 15 µL of respective buffers (kit components). Finally, 75 µL of $NAD^+$/NADH reaction mixture ($NAD^+$/NADH recycling enzyme mixture and sensor buffer, kit components) were added and the resulting mixtures incubated for 1 h at room temperature. Fluorescence intensities (ex. 540 nm; em. 590 nm) were recorded using a Perkin Elmer Wallac 1420 Victor2 Microplate Reader. Values were derived from the standard curve of each experiment and compared to the data obtained by mass spectrometry-based metabolomics, to calculate total and mitochondrial NADH concentrations.

***Quantitative metabolomics:*** In a typical experiment 1.5 million cells were used for total extracts and 15 million cells for mitochondrial extracts. Cells were harvested and the supernatant removed to generate the corresponding cell pellets. Subsequently, pellets were dried and dry pellets were supplemented with 300 µl methanol, vortexed 5 min and centrifuged (10 min at 15000 g, 4 °C). Then, the upper phase of the supernatant was split into two parts: 150 µL were used for a gas chromatography coupled by mass spectrometry (GC/MS) experiment in microtubes and the remaining 150 µL were used for Ultra High Pressure Liquid Chromatography coupled by Mass Spectrometry (UHPLC/MS). For the GC-MS aliquots, supernatants were completely evaporated from the sample. 50 µL of methoxyamine (20 mg/mL in pyridine) were added to the dried extracts, then stored at room temperature in the dark for 16 h. The following day, 80 µL of MSTF (A-Methyl-N-(trimethylsilyl) trifluoroacetamide) were added and final derivatization occurred at 40 °C for 30 min. Samples were then transferred into vials and directly injected for GC-MS analysis. For the UHPLC-MS aliquots, 150 µL were dried in microtubes at 40 °C in a pneumatically-assisted concentrator (Techne DB3, *Staffordshire, UK*). The dried UHPLC-MS extracts were solubilized with 200 µL of MilliQ water. Aliquots for analysis were transferred into LC vials and injected into UHPLC-MS or kept at -80 °C until injection.

*Widely-targeted analysis of intracellular metabolites gas chromatography (GC) coupled to a triple quadrupole (QQQ) mass spectrometer.* GC-MS/MS method was performed on a 7890A gas chromatography (Agilent Technologies, Waldbronn, Germany) coupled to a triple quadrupole 7000C (Agilent Technologies, Waldbronn, Germany) equipped with a High sensitivity electronic impact source (EI) operating in positive mode[68]. Peak detection and integration of the analytes were performed using the Agilent Mass Hunter quantitative software (B.07.01).

*Targeted analysis of nucleotides and cofactors by ion pairing ultra-high performance liquid chromatography (UHPLC) coupled to a Triple Quadrupole (QQQ) mass spectrometer.* Targeted analysis was performed on a RRLC 1290 system (Agilent Technologies, Waldbronn, Germany) coupled to a Triple Quadrupole 6470 (Agilent Technologies) equipped with an electrospray source operating in both negative and positive modes. Gas temperature was set to

350 °C with a gas flow of 12 L/min. Capillary voltage was set to 5 kV in positive mode and 4.5 kV in negative mode. 10 µL of sample were injected on a Column Zorbax Eclipse XDB-C18 (100 mm × 2.1 mm particle size 1.8 µm) from Agilent technologies, protected by a guard column XDB-C18 (5 mm × 2.1 mm particle size 1.8 µm) and heated at 40 °C by a pelletier oven. The gradient mobile phase consisted of water with 2 mM of dibutylamine acetate concentrate (DBAA) (A) and acetonitrile (B). Flow rate was set to 0.4 mL/min and an initial gradient of 90% phase A and 10% phase B, which was maintained for 3 min. Molecules were then eluted using a gradient from 10% to 95% phase B over 1 min. The column was washed using 95% mobile phase B for 2 minutes and equilibrated using 10% mobile phase B for 1 min and the autosampler was kept at 4 °C. Scan mode used was the MRM for biological samples. Peak detection and integration of the analytes were performed using the Agilent Mass Hunter quantitative software (B.10.1).

*Pseudo-targeted analysis of intracellular metabolites by ultra-high performance liquid chromatography (UHPLC) coupled to a Q-Exactive mass spectrometer. Reversed phase acetonitrile method.* The profiling experiment was performed with a Dionex Ultimate 3000 UHPLC system (Thermo Scientific) coupled to a Q-Exactive (Thermo Scientific) equipped with an electrospray source operating in both positive and negative modes and full scan mode from 100 to 1200 m/z. The Q-Exactive parameters were: sheath gas flow rate 55 au, auxiliary gas flow rate 15 au, spray voltage 3.3 kV, capillary temperature 300 °C, S-Lens RF level 55 V. The mass spectrometer was calibrated with sodium acetate solution dedicated to low mass calibration. 10 µL of sample were injected on a SB-Aq column (100 mm × 2.1 mm particle size 1.8 µm) from Agilent Technologies, protected by a guard column XDB-C18 (5 mm × 2.1 mm particle size 1.8 µm) and heated at 40 °C by a pelletier oven. The gradient mobile phase consisted of water with 0.2% acetic acid (A) and acetonitrile (B). The flow rate was set to 0.3 mL/ min. The initial condition was 98% phase A and 2% phase B. Molecules were then eluted using a gradient from 2% to 95% phase B for 22 min. The column was washed using 95% mobile phase B for 2 for min and equilibrated using 2% mobile phase B for 4 min. The autosampler was kept at 4 °C. Peak detection and integration were performed using the Thermo Xcalibur quantitative software (2.1.) [68].

***Energy calculation of copper(II)-catalyzed hydride transfer to $H_2O_2$ from NADH:*** The UBHLYP functional[61,62] was used associated with the SVP basis set[69,70] and the SMD solvation model[71,72] to represent an adequate method to describe the $[Cu(H_2O)_6]^{2+}$ species[65]. Thus, all structures (minima and transition states) were optimized using the Gaussian 16 set of programs at the UBHLYP/SVP level for all atoms (doublet spin state). The SMD solvation model (water) was applied during the optimization process. Thermal correction to the Gibbs free energy was computed at 310.15 K. Single points at the UMP2/SVP level were performed. The results presented are $\Delta G_{298}$ in kcal/mol. 1-Methyl-1,4-dihydronicotinamide was chosen as a NADH model to study the copper(II)-catalyzed hydride transfer to $H_2O_2$.

***Cytokine quantification:*** Cytokines levels were measured in cell culture supernatants using the V-Plex pro-inflammatory panel (MSD, Rockville, MD, US, #K15049D-1). The kit was run according to the manufacturer's protocol and the chemiluminescence signal was measured on a Sector Imager 2400 (MSD).

***RNA-seq:*** *Human samples:* RNAs were extracted using the RNeasy mini kit (QIAGEN, 74104). *Mouse samples:* half of the right lobe was homogenized in 1 mL of RA1 buffer from the NucleoSpin RNA kit (Macherey Nagel, 740955.250) containing 20 mM of tris(2-carboxyethyl)phosphine (TCEP). Total RNAs in the tissue homogenate were extracted with the NucleoSpin RNA kit. RNAs were eluted with 60 µL of water. RNA sequencing libraries were prepared from 1 μg total RNA using the Illumina TruSeq Stranded mRNA library preparation kit (Illumina, 20020594), which allows strand-specific sequencing. A first step of polyA selection using magnetic beads was performed to allow sequencing of polyadenylated transcripts. After fragmentation, cDNA synthesis was performed and resulting fragments were used for dA-tailing followed by ligation of TruSeq indexed adapters (Illumina, 20020492). Subsequently, polymerase chain reaction amplification was performed to generate the final barcoded cDNA libraries. Sequencing was carried out on a NovaSeq 6000 instrument from Illumina based on a 2× 100 cycle mode (paired-end reads, 100 bases). Raw sequencing reads were first checked for quality with Fastqc (0.11.8) and trimmed for adapter sequences with the trimGalore (0.6.2) software. Trimmed reads were then aligned on the human hg38 or mouse mm10 reference genome using the STAR mapper (2.6.1b), up to the generation of a raw count table per gene (GENCODE annotation v29). The bioinformatics pipelines used for these tasks are available online (rawqc v2.1.0: https://github.com/bioinfo-pf-curie/raw-qc, RNA-seq v3.1.4: https://github.com/bioinfo-pf-curie/RNA-seq). The downstream analysis was then restricted to protein-coding genes. Data from[73] were converted into bulk by keeping cells annotated as macrophages and then summing the counts for each sample. Counts data from[74] were downloaded from GEO under accession number GSE73502. Raw data from[75,76] were downloaded from the NCBI Short Read Archive under records PRJNA528433 and PRJNA290995 and processed as described above. Counts were normalized using TMM normalization from edgeR (v 3.30.3)[77]. Differential expression was assessed with the limma voom framework (v 3.44.3)[78]. The intra-donor correlation was controlled by using the duplicateCorrelation from limma. Genes with an adjusted $p$-value<0.05 were labeled significant. Enrichment analysis from differentially expressed genes has been performed using the enrichGO function from clusterProfilter package v3.16.1.

***LPS-induced severe inflammation mouse model:*** Survival assessment using the lipopolysaccharide (LPS) mouse model was conducted at Fidelta Ltd according to 2010/63/EU and National legislation regulating the use of laboratory animals in scientific research and for other purposes (Official Gazette 55/13). An Institutional Committee on Animal Research Ethics (CARE-Zg) oversaw that animal-related procedures were not compromising the animal welfare. LPS (Sigma-Aldrich, L2630, 20 mg/kg) was injected intraperitoneally to male BALB/c mice (8 weeks old). LCC-12 (0.3 mg/kg, IP, $n$=10) or vehicle (0.9% NaCl, 10 mL/kg, IP, $n$=10) were injected 2 h prior LPS challenge, then 24 h, 48 h, 72 h and 96 h post challenge. Dexamethasone (10 mg/kg, PO, $n$=10) was given 1 h prior LPS challenge. Incidence of mortality was monitored every 4 h up to 48 h, then twice daily. The cytometry work and ICP-MS work using the LPS mouse models were performed in accordance with French laws concerning animal experimentation (#2021072216346511) and approved by Institutional Animal Care and Use Committee (C2EA-47). Experiments were performed on 8-week-old male BALB/c mice and 5-week-old male SWISS mice. Male littermates were randomly assigned to experimental groups. The experimental endotoxemic model was induced by intraperitoneal (IP) injection of LPS (5 mg/kg in BALB/c mice (*Escherichia coli* O111:B4, Sigma-Aldrich, L2630) or 20 mg/kg in SWISS mice (*Escherichia coli* O55:B5, Sigma-Aldrich, L2880)). All the animals were resuscitated with 30 mL/kg body weight of saline administered subcutaneously 6 h post LPS administration. Mice were intra-peritoneally injected with LCC-12 (0.3 mg/kg 2 h before LPS challenge for BALB/c mice or 6 h after for SWISS mice). Mice were sacrificed 22 h post-LPS challenge. The temperature was measured at 0 h, 6 h and 22 h post-LPS challenge. *Flow cytometry:* After euthanasia of mice at 22 h post LPS challenge, the organs were perfused with PBS/EDTA (1 mL/g, 2mM, pH 7.4) and 10 mL of 1× PBS were injected in the peritoneum. The peritoneal liquid was then collected and centrifugated for 5 minutes at 1,500 rpm. The pellet was resuspended in RPMI medium containing 2% fetal calf

serum (Dutscher, S181H-100). The peritoneal liquid was then washed in 96-well plates at 2,000 rpm for 2 min, and pellets were suspended in complete RPMI. All samples were incubated or not for 2 h in brefeldin A (BFA, 5 ng/mL, 00-4506-51, ThermoFischer) before surface staining with Fixable Viability Dye eFluor 780 (Thermofisher, 65-0865-14) followed by fluorochrome-conjugated antibodies (35 min at 4°C). Samples incubated with BFA were fixed (Foxp3/ Transcription factor Fix/Perm 4X, TONBO Biosciences, 44931S) for 20 min at 4°C and permeabilized (Flow Cytometry Perm Buffer 10X, TONBO Biosciences, TNB-1213-L150) before intracellular staining. The antibodies used were as follows: CD11b-Pacific Blue (BioLegend, 101224), CD40-APC (BioLegend, 124612), CD45-BV510 (BioLegend, 103138), CD71-BV711 (BD Biosciences, 740667), CD86-PE (Biolegend, 105007), CD170-PEeFluor 610 (eBioscience, 61-1702-80), CD206-BV650 (BioLegend, 141723), F4/80-BV605 (BioLegend, 123133), I-A/I-E-AF700 (BioLegend, 107622), Ly6C-PerCP/Cy5.5 (BioLegend, 128012) and Ly6G-PE/CY7 (BioLegend, 127618). For intracellular staining, NOS2-APC (eBioscience, 17-5920-82) was used. Small peritoneal macrophages (SPM) correspond to CD45$^+$/IA-IE$^+$/CD11b$^+$/F4/80$^{int}$/SiglecF$^-$ cells. After washing in PBS or in Perm Buffer, data were acquired using a LSR Fortessa flow cytometer (BD Biosciences, France) and analyzed with FlowJo.10 software.
*Inductively coupled plasma mass spectrometry (ICP-MS):* ICP-MS experiments were conducted as described in the ICP-MS paragraph on peritoneum (omentum) tissues and SPM. Tissue-specific data were normalized against dry weight. The sorting of SPM was done using the following antibodies: CD11b-Pacific Blue (BioLegend, 101224), F4/80-PE (TONBO, TNB50-4801-U100), Ly6C-PerCP/Cy5.5 (BioLegend, 128012) and Ly6G-AF647 (BioLegend, 127610).

***Cecal ligation and puncture mouse model:*** All animal-related research is conducted in accordance with 2010/63/EU and National legislation regulating the use of laboratory animals in scientific research and for other purposes (Official Gazette 55/13). An Institutional Committee on Animal Research Ethics (CEEA - 047) oversees that animal-related procedures are not compromising the animal welfare. 9-week-old male BALB/c mice were used for these experiments. Animals were anesthetized by isoflurane (Forene). After abdominal incision, the cecum was ligated, punctured with a gauge needle (25G), and a small amount of fecal matter was released. After the cecum was returned to the abdomen, the abdominal cavity was closed in two layers and the mice were resuscitated with 30 mL/kg body weight of saline (0.9% NaCl) administered subcutaneously. For the sham group, after abdominal incision, the cecum was manipulated but was neither ligated nor punctured. After the cecum was returned to the abdomen, the abdominal cavity was closed in two layers and the mice were resuscitated with 30 mL/kg body weight of saline administered subcutaneously. LCC-12 (0.3 mg/kg, IP) was administered at 0.3 mg/kg dose 4 h, 24 h, 48 h, 72 h and 96 h following CLP creation. Mortality incidence was monitored every 2 h up to 120 h (except from 10 pm to 6am) post CLP creation. Dexamethasone was administered intraperitoneally at 1 mg/kg dose 5-minutes prior CLP creation.

***SARS-CoV-2 mouse model:*** All experiments involving SARS-CoV-2 were performed within the biosafety level 3 facility of the Institut Pasteur de Lille, after validation of the protocols by the local committee for the evaluation of the biological risks and complied with current national and institutional regulations and ethical guidelines (Institut Pasteur de Lille/B59-350009). The experimental protocols using animals were approved by the institutional ethical committee "Comité d'Ethique en Experimentation Animale (CEEA) 75, Nord-Pas-de-Calais". The animal study was authorized by the "Education, Research and Innovation Ministry" under registration number APAFIS#25517-2020052608325772v3. 8-week-old male K18-human ACE2 expressing C57BL/6 mice (B6.Cg-Tg(K18-hACE2)2Prlmn/J) were purchased from Jackson Laboratory. The mice were anesthetized by IP injection of ketamine (100 mg/kg) and xylazine (10 mg/kg) and then intranasally infected with 50 µl of DMEM containing 5x10$^{2}$ $TCID_{50}$ of hCoV-19_IPL_France strain of SARS-CoV-2 (NCBI MW575140). LCC-12 was inoculated intranasally (0.5 mg/ml, 50 µL) 6 h, 24 h and 48 h post-infection. Mice were sacrificed at day 4 post-infection.

*RT-qPCR:* Half of the right lobe was homogenized in 1 mL of RA1 buffer from the NucleoSpin RNA kit containing 20 mM of TCEP. Total RNAs in the tissue homogenate were extracted with NucleoSpin RNA from Macherey Nagel. RNAs were eluted with 60 µL of water. RNA was reverse-transcribed with the High-Capacity cDNA Archive Kit (Life Technologies, USA). The resulting cDNA was amplified using SYBR Green-based real-time PCR and the QuantStudi 12K Flex Real-Time PCR Systems (Applied Biosystem, USA) following the manufacturer's protocol. Relative quantifications were performed using the gene coding for glyceraldehyde 3-phosphate dehydrogenase (Gapdh). Specific primers were designed using Primer Express software (Applied Biosystems, Villebon-sur-Yvette, France) and ordered from Eurofins Scientifics (Ebersberg, Germany). Relative mRNA levels ($2^{-\Delta\Delta Ct}$) were determined by comparing (a) the PCR cycle thresholds (Ct) for the gene of interest and the house keeping gene (ΔCt) and (b) ΔCt values for treated and control groups (ΔΔCt). Data were normalized against expression of the *GAPDH* gene and are expressed as a fold-increase over the mean gene expression level in mock-treated mice. Primer sequences used are tabulated below:

| Gene | Accession Number | Forward primer 5′-3′ | Reverse primer 5′-3′ |
|---|---|---|---|
| *Gapdh* | NM_001289726 | GCAAAGTGGAGATTGTTGCCA | GCCTTGACTGTGCCGTTGA |
| *Ifng* | NM_008337.4 | CAACAGCAAGGCGAAAAAG | GTGGACCACTCGGATGAGCT |
| *Il6* | NM_031168.2 | CAACCACGGCCTTCCCTACT | CCACGATTTCCCAGAGAACATG |
| *Isg15* | NM_015783.3 | GGCCACAGCAACATCTATGAGG | CTCGAAGCTCAGCCAGAACTG |
| *Mx1* | NM_010846.1 | TGCAGAGGTCAGCAGGACATC | GGCAGTTTGGACCATCTCTGAA |
| *Stat1* | NM_019663.3 | GCTGCCTATGATGTCTCGTTTG | TTCCGTATGTTGTGCTGCAAC |

*CD44 immunohistochemical staining:* Lung tissues were fixed in 1× PBS with 4% formaldehyde for 7 d, rinsed in 1× PBS, transferred into ethanol (70%) and then processed into paraffin-embedded tissue blocks and 3 µm tissue sections were cut. Slides were heated at 60 °C for 1 h, deparafinized in xylene and rehydrated in ethanol baths. Target retrieval (pH 9.0) was then performed in citrate buffer (Agilent Dako, S2367) at 95 °C for 30 min. This step was followed by successive incubations in peroxidase blocking solution for 10 min (Dako, S2023), serum-free protein blocking solution for 10 min (Dako, X0909) and CD44 antibody for 60 min (Abcam, ab189524, 1:4000). Target detection was done by incubation in a secondary antibody conjugated with HRP (Dako, K4003), then revealed using 3-Amino-9-ethylcarbazole (AEC). Slides were then stained with hematoxylin (Sigma-Aldrich, HHS32-1L). Analysis was performed using QuPath software version 3.0. Tissue area was calculated using a classifier separating cells from blank space. This way, blank space within vessels and alveoli was removed and only lung tissue containing cells was kept. Cell detection was performed by adjusting parameters including cell size, fragmentation or hematoxylin detection threshold. Positive detection was then performed using AEC intensity. Cell density was calculated using the following ratio: positive cell count/tissue area.

***Software for illustrations:*** Illustrations were created using Adobe Illustrator and biorender.com.

***Quantification and statistical analysis:*** Results are presented as mean values ± standard error of the mean (SEM) or standard deviation (SD) as indicated. Box plots are plotted with the median and whiskers of lowest and highest values. PRISM 8 software was used to calculate *p*-values using a Mann-Whitney test, Student's T-test, Kruskal-Wallis test with Dunn's post-test, 2-way ANOVA or Mantel-Cox log-rank test as indicated. PRISM 8 software or R programming language was used to generate graphical representations of quantifications unless stated otherwise. Sample sizes (*n*) are indicated in the figure legends.
